# Reverse molecular pharmacology identifies the non-canonical axis of IRAK as a chemoresistance factor in neuroblastoma

**DOI:** 10.64898/2026.01.09.697473

**Authors:** M Le Grand, C Buxbaum, M Kerhervé, F Gaucher, Á Martínez-Rubio, M Taha, K Muller, B Mouysset, A Bomane, S Letard, T.W Failes, G.M Arndt, S Chebbi, E Labaronne, F.M.G Cavalli, N André, L Broutier, Y Shaked, E Pasquier

## Abstract

Owing to chemoresistance, the prognosis of relapsed neuroblastoma is dismal with less than 10% of patients surviving after 5 years. We developed a reverse molecular pharmacology approach that is based on high-throughput drug screening coupled with chemo-informatic and transcriptomic analyses. This led to the identification of IRAK1 as a key chemoresistance factor in neuroblastoma. By performing functional and pharmacological drug combination screens targeting IRAK1, we revealed a synergy between IRAK1 inhibition/silencing and BET, EGFR and mTOR inhibitors as well as microtubule-targeting agents. The synergistic combination of microtubule-targeting agent, vincristine and IRAK inhibitors was then confirmed in tumor spheroids, patient-derived tumoroids and a syngeneic orthotopic mouse model. Mechanistically, IRAK inhibition potentiated the pro-apoptotic and cell cycle arrest properties of vincristine *via* a pathway involving the PIDDosome complex rather than its canonical MyDDosome axis. Altogether, this study represents a proof-of-concept of our reverse molecular pharmacology approach to quickly develop biology-guided drug combinations, that could be applied to any other human diseases.

## INTRODUCTION

With the emergence of precision oncology strategy, the use of targeted therapies to benefit patients whose tumors exhibit specific molecular traits has exploded ^1^. However, it is now well-established that most effective drugs in cancer, including the targeted therapies act on multiple rather than single targets ^2,3^. This is reflected with the ever-increasing number of chemo-informatics databases that collect drug information such as chemical structures and target engagement, demonstrating the high level of drug polypharmacology ^4–6^. Thus, over the last decade, the concept of drug poly-pharmacology has arisen as a unique opportunity to identify key vulnerabilities in specific diseases, opening major therapeutic avenues ^7–9^. This is perfectly illustrated by our previous study where we established a biology-guided drug combination screening method. This allowed us to identify the dual inhibition of AURKA and BET proteins as a promising strategy in glioblastoma, providing sound foundations for future clinical trials ^10^.

Neuroblastoma (NB) is an enigmatic, multifaceted tumor of the peripheral nervous system that represents the most common extracranial solid tumor in children under five years of age ^11^. This neuroendocrine tumor originates most frequently in the adrenal gland, but it can also develop in the neck, chest, abdomen, or pelvis. Based on its cellular and biological heterogeneity, NB behavior can range from low-risk tumors with a tendency toward spontaneous regression, to high-risk ones with extensive growth, early metastasis and a poor prognosis ^12,13^. High-risk NB represent almost 50% of all cases and have a poor prognosis with a 5-year survival rate below 40%. Half of these patients will relapse despite intensive multimodal treatment based on four therapeutic phases: an intensive induction chemotherapy, a local control by surgery and/or radiation, a consolidation phase with high dose chemotherapy and autologous transplant and a maintenance phase with anti-GD2 followed by differentiating therapy with polyamine inhibitor DFMO ^14,15^. Owing to acquired resistance mechanisms, the prognosis at relapse is even more dismal with less than 10% of patients surviving after 5 years ^12^.

The interleukin-1 receptor–associated kinase (IRAK) family comprises four isoforms that serve as key mediators of toll-like receptor (TLR) signaling, controlling diverse cellular processes such as inflammation and innate immunity ^13^. Several studies have demonstrated their oncogenic role by enhancing tumor angiogenesis, promoting tumor growth and contributing to chemo- and radio-resistance ^17–21^. Few IRAK inhibitors are currently in preclinical and clinical development for human cancers to assess whether their potential will translate to clinical efficacy ^22^. However, the roles of the IRAK family members in NB biology and chemoresistance have not yet been explored.

In this study, we developed a reverse molecular pharmacology approach to unveil chemoresistance factors in NB. By conducting functional and pharmacological high-throughput screening followed by chemo- and bio-informatic analyses we identified IRAK1 as a regulator of NB chemoresistance. We further explored the biological and therapeutic roles of IRAK in NB. bm

## RESULTS

### Reverse molecular pharmacology approach identified seven actionable vulnerabilities involved in an acquired resistance model of neuroblastoma

Chemotherapy is a major component of standard treatment for high-risk NB patients^15^. However, drug resistance frequently develops, causing disease relapse. To uncover new actionable vulnerabilities that could overcome chemoresistance, we developed an approach called reverse molecular pharmacology based on high-throughput drug screening combined with chemo- and bio-informatic analyses. We used a high-risk NB cell line (BE/VCR10) that has acquired resistance to vincristine (a common cytotoxic drug used in clinic) after chronic exposure (IC_50_ of 28 ± 7 nM and 2,767 ± 361 nM for parental BE(2)-C and BE/VCR10, respectively; Supp. 1A-B). The resistant cell line was used to screen 3 compound libraries, containing ∼2,800 different already-approved drugs and pharmacologically active molecules (Fig. 1A; Supp Table 1), as monotherapies and in combination with vincristine. Seventy-six compounds were able to inhibit NB cell viability by at least 75% including 14 chemotherapy agents, 13 kinase inhibitors and 7 anti-helminthic drugs (Supp. 2C and Supp. Table 2 for details). Moreover, hit compounds that could re-sensitize BE/VCR10 cells to vincristine by at least 25% were considered as potential chemosensitizers. Ninety-one compounds met this criterium and were used in a secondary screen. Sixty-six compounds were further validated (Fig. 1B and Supp. Table 3). As our method aims at identifying actionable vulnerabilities involved in chemoresistance, we next focused on these 66 compounds. By mining chemo-informatics databases (Fig. 1A), a list of 195 putative therapeutic targets, with an average of 8.3 targets per hit compound was generated (Supp. Table 4). KEGG pathway enrichment analysis indicated that the 195 selected targets were enriched in 4 main pathways: neuroactive ligand-receptor interaction, calcium signaling pathway, serotoninergic synapse and MAPK signaling pathway (Fig. 1C, and Supp. Table 4 for details). Exploiting the R2 online platform, we then conducted gene expression profiling using two independent cohorts of NB patients (#GSE45547; #GSE 49710; Fig. 1A). Amongst the 195 putative targets, we identified 7 genes, whose high expression significantly correlates with poor prognosis in both cohorts: three kinases (*IRAK1, RET* and *STK33*), one methyltransferase (*PRMT1*), one isomerase (*EBP*), one progesterone binding protein (*PGRMC1*) and one serine/threonine protein phosphatase (*PPP1CA*) (Fig. 1D). Seven compounds from the original drug screening target these genes including gefitinib or mometasone (Fig. 1D). The therapeutic potential of specifically targeting these candidates was then assessed using the Drug–Gene Interaction database ^23^. Pharmacological compounds were referenced for all seven targets. Moreover, the expression of the selected genes was confirmed in a panel of 12 NB models harboring different molecular features (Fig. 1E).

**Figure 1.**
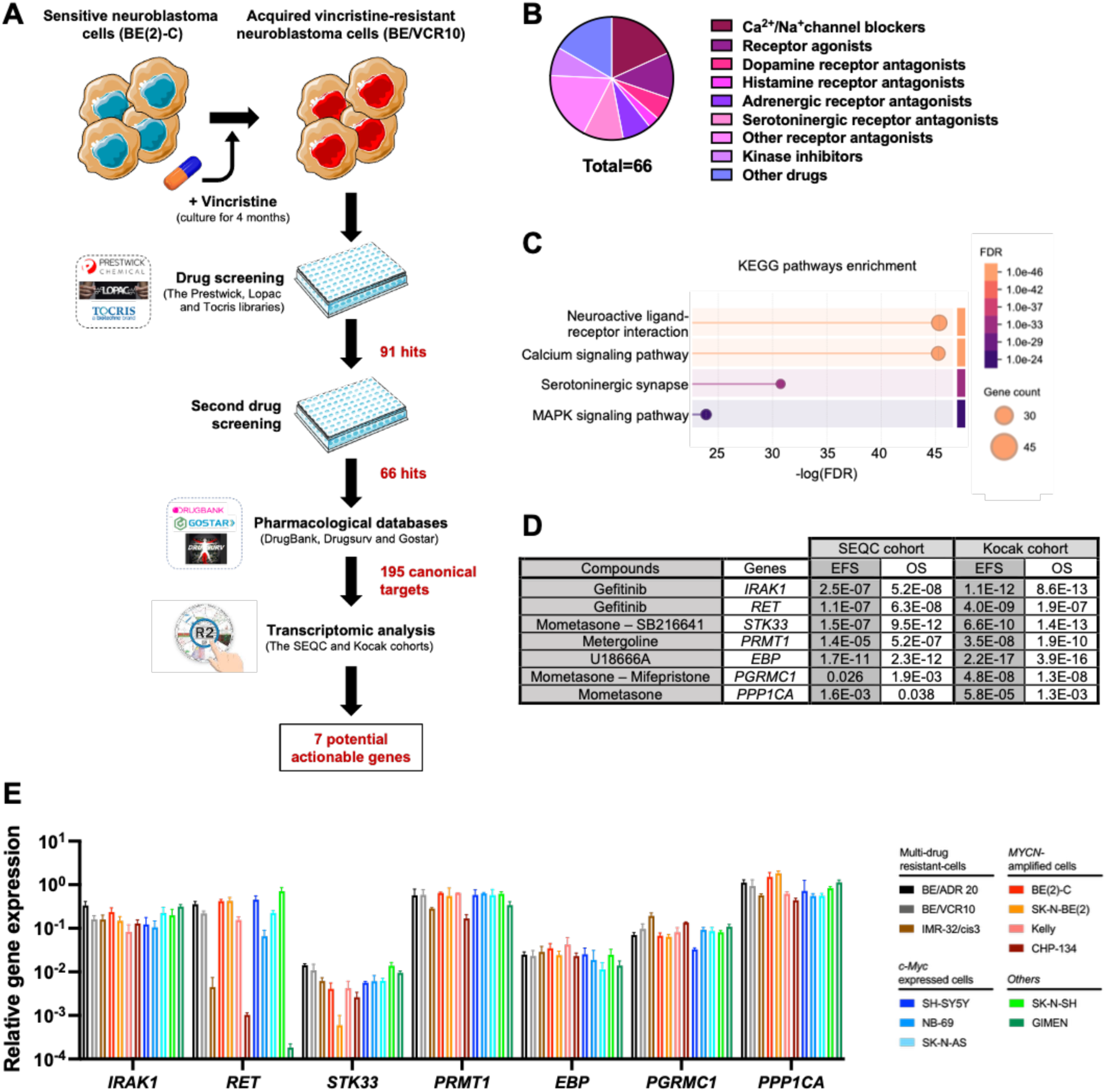
High-throughput drug screening combined with chemo- and bio-informatic analyses identified seven vulnerabilities in acquired resistance neuroblastoma model. (**A**) Workflow of our reverse molecular pharmacology approach. (**B**) Sixty-six compounds were defined as NB chemo-sensitizers (>25% increase in vincristine sensitivity) and are represented as a donut diagram and classified by pharmacological classes. (**C**) Top 4 enriched KEGG pathways ranked by -logFDR based on the 195 targets of the chemo-sensitizers identified using STRING network analysis (https://string-db.org). (**D**) Univariate analysis of event free survival (EFS) and overall survival (OS) of NB patients from the SEQC cohort (n=498) and the Kocak cohort (n=609) regarding the seven identified targetable vulnerabilities. Median cut-off of the gene expression was used. (**E**) Gene expression in a panel of 12 NB cell lines by qRT-PCR using *YWHAZ* as housekeeping gene. *Bars*, mean of at least 4 independent experiments; *Error bars*, S.D.

### *IRAK1* regulates cell viability, tumor spheroid growth and vincristine sensitivity in neuroblastoma models

To explore the physiological relevance of targeting the seven identified genes, we performed a transient knockdown of each target using small interfering RNAs (siRNAs) in the multi-resistant BE/VCR10 cell line and the parental cell line BE(2)-C (Fig. 2A). First, our data indicated that the viability of BE/VCR10 cells was significantly reduced following the downregulation of 5 out of 7 targets, including *PPP1CA*, *IRAK1*, *RET*, *STK33* and *PRMT1* (Fig. 2AB grey line, *p* < 0.05). These results were confirmed in the BE(2)-C model with the downregulation of *IRAK1*, *RET*, and *PRMT1*, while a significant decrease in the BE(2)-C cell viability was also observed following *EBP* knockdown (Fig. 2B, red line, *p* < 0.05). In order to confirm these data, we employed BE/VCR10 3D tumor spheroids, which better mimic tumor characteristics. A transient knockdown of *IRAK1*, *RET*, *STK33* and *PRMT1* resulted in a decrease in tumor spheroid viability, observed at day 8: 32 ± 9%, 26 ± 8%, 27 ± 12% and 23 ± 8% for the downregulation of *IRAK1*, *RET*, *STK33* and *PRMT1* in comparison to control, respectively (Fig. 2B, black line, *p* < 0.05). Since only the downregulation of *IRAK1*, *RET* and *PRMT1* led to a significant decrease in all tested models (Fig. 2B), we checked the impact of knocking down their gene expression in the BE(2)-C and SK-N-SH spheroid models (Fig. 2C). Our results indicated that a significant decrease in spheroid viability in both models was only observed with the downregulation of *IRAK1* and *RET*. Indeed, *IRAK1* silencing resulted in a drop of 51 ± 8% and 30 ± 6% (Fig. 2BD, *p* ≤ 0.01) in BE(2)-C and SK-N-SH spheroid viability, respectively, while *RET* knockdown led to a decrease of 21 ± 7% and 22 ± 3% (Fig. 2D, *p* ≤ 0.05) in the two models. Then, we assessed the interest of targeting these 7 candidate genes to overcome chemoresistance. The downregulation of *IRAK1* led to an increase in vincristine sensitivity in BE(2)-C cells (Fig. 2E). To confirm the impact of *IRAK1* silencing, a transient knockdown was performed in 5 different 3D spheroid models of NB harboring different molecular characteristics (Fig. 2F-G). Our results showed that the downregulation of *IRAK1* significantly sensitized the BE/VCR10, BE(2)-C, SK-N-BE(2) and SH-SY5Y spheroids to vincristine, while a similar trend was observed in the SK-N-SH spheroids (Fig. 2H). Altogether, by applying our reverse molecular pharmacology approach, we identified gefitinib as a NB chemo-sensitizer and uncovered that IRAK1, one of its targets is a key factor in cell and tumor spheroid viability and vincristine sensitivity in NB.

**Figure 2.**
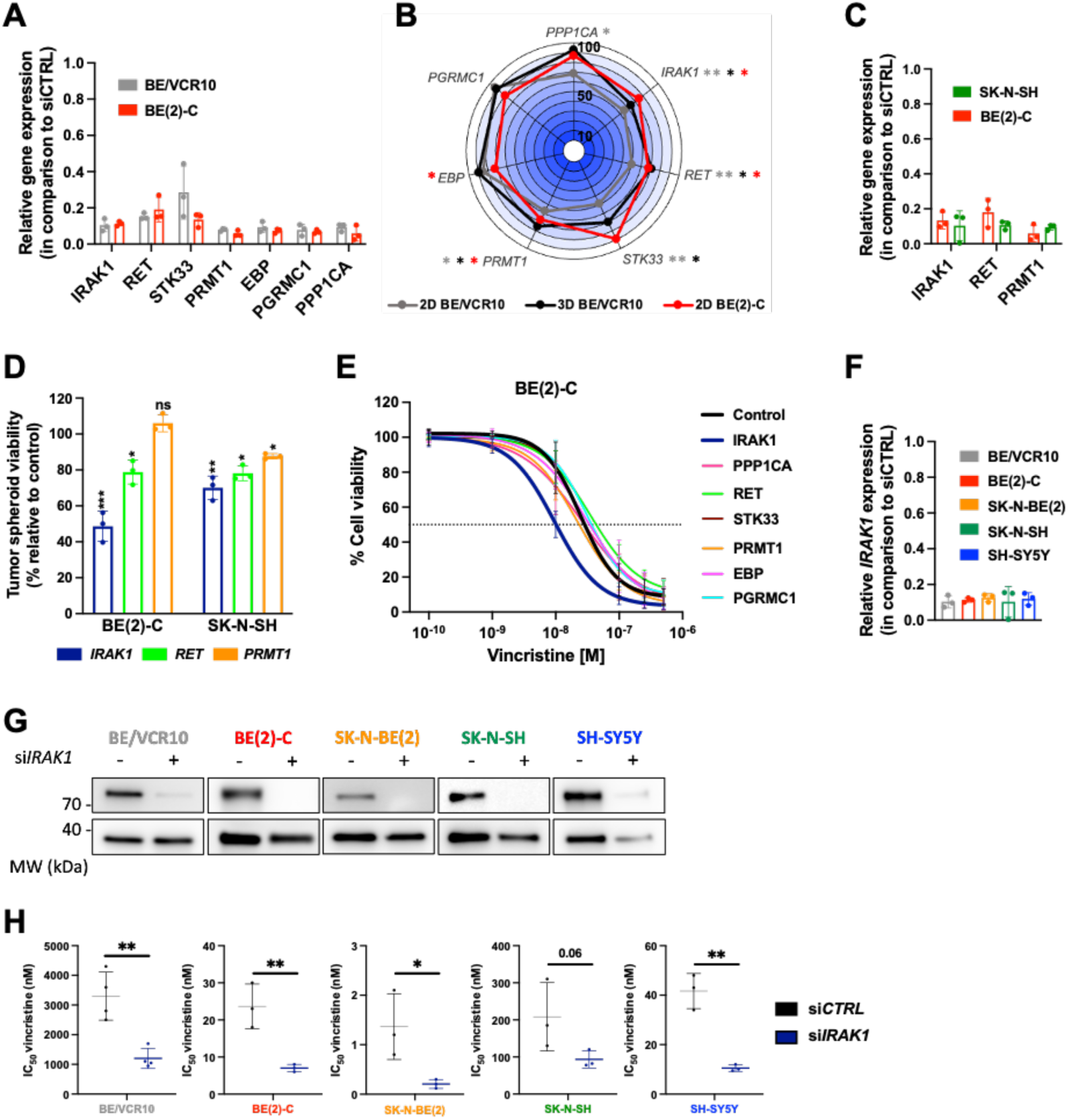
*IRAK1* regulates cell viability, tumor spheroid growth and vincristine sensitivity in neuroblastoma cells. (**A**) Relative gene expression following 48h transfection of BE/VCR10 or BE(2)-C cells with negative control siRNA and siRNA sequences targeting the 7 target hits, as evaluated by qRT-PCR using *YWHAZ* as housekeeping gene. (**B**) Polar plot representing the viability of BE/VCR10 (grey line) and BE(2)-C (red line) cells and BE/VCR10 spheroids (black line) following the transient downregulation of the 7 gene hits. Polar plot was made up of 10 data rings, each radial point representing a ten percent increment of cell viability on a scale from 0 (inner radial point) to 100 (outer radial point). *Dots*, mean of at least 3 independent experiments; * *p <* 0.05; *\*\* p* ≤ 0.01. (**C**) Relative gene expression following 48h transfection of SK-N-SH or BE(2)-C cells with negative control siRNA and siRNA sequences targeting *IRAK1*, *RET* or *PRMT1*, as evaluated by qRT-PCR using *YWHAZ* as housekeeping gene. (**D**) Histograms representing BE(2)-C and SK-N-SH tumor spheroid viability at day 8 following siRNA transfection targeting *IRAK1*, *RET* and *PRMT1*. Values are the average of at least three independent experiments ± S.D; ns, p>0.05; *, *p <* 0.05; *\*\*, p* ≤ 0.01. (**E**) Dose–response curves of BE(2)-C cells either transfected with negative control or siRNA targeting one of the 7 gene hits, treated with vincristine for 72 hours. (**F**) Relative gene expression following 48h transfection in a panel of 5 NB cells with negative control siRNA and siRNA sequences targeting *IRAK1*, as evaluated by qRT-PCR using *YWHAZ* as housekeeping gene. (**G**) Representative western blot showing IRAK1 protein expression following 72h siRNA transfection in BE/VCR10, BE(2)-C, SK-N-BE(2), SK-N-SH and SH-SY5Y, using GAPDH as loading control. (**H**) IC_50_ values of vincristine calculated after 7-days treatment in BE/VCR10, BE(2)-C, SK-N-BE(2), SK-N-SH and SH-SY5Y 3D spheroid models either transfected with negative control or siRNA targeting *IRAK1*. *Bars*, mean of at least 3 independent experiments; *Error bars*, S.D; *, *p <* 0.05; *\*\*, p* ≤ 0.01.

### IRAK1 is a clinically-relevant therapeutic target in neuroblastoma

The IRAK family comprises 4 different isoforms: IRAK1, 2, 3 and 4 ^24^. To investigate the clinical significance of *IRAK* isoform expression levels, we used publically available microarray data from two large cohorts of NB patients (#GSE45547; #GSE 49710) and the Genotype-Tissue Expression (GTeX) database gathering normal tissue records. All *IRAK* isoforms were upregulated at the gene level in NB patients as compared to normal adrenal gland tissue in both cohorts (fold change > 1.5; *p* < 0.001; Fig. 3A). Next, high expression of *IRAK1* was significantly associated with shorter event-free and overall survival (EFS and OS; *p* < 0.001; Bonferroni-corrected, Log-rank test) in an independent cohort of NB patients (n = 498; SEQC cohort), while the opposite trend was observed with *IRAK2*, *IRAK3* and *IRAK4* (Fig. 3B and Supp. 3A-B). Moreover, *IRAK1* gene expression was significantly elevated in *MYCN*-amplified NB tissue in comparison to non-*MYCN*-amplified NB tissue and in INSS stage 4 disease relative to earlier stages (*p* < 0.001), while the gene expression levels of other isoforms dropped in patients with high-risk disease (*i.e* INSS stage 4 and *MYCN*-amplified tissue; Fig. 3C-D). We finally performed a multivariate analysis to determine whether the prognostic significance of *IRAK1* expression was independent of established prognostic indicators for NB. Age at diagnosis (< 18 months *vs.* > 18 months), tumor stage (1 *vs.* 2,3,4 or 4s), *MYCN* status (amplified *vs.* single copy), and *IRAK* isoform gene expression were tested in a Cox proportional hazards regression model. Our results demonstrated that only high *IRAK1* expression retained strong independent prognostic significance for EFS and OS in NB patients (Supp. Tables 5 and 6). Collectively, these results indicate that IRAK1 could be a clinically-relevant target in NB.

**Figure 3.**
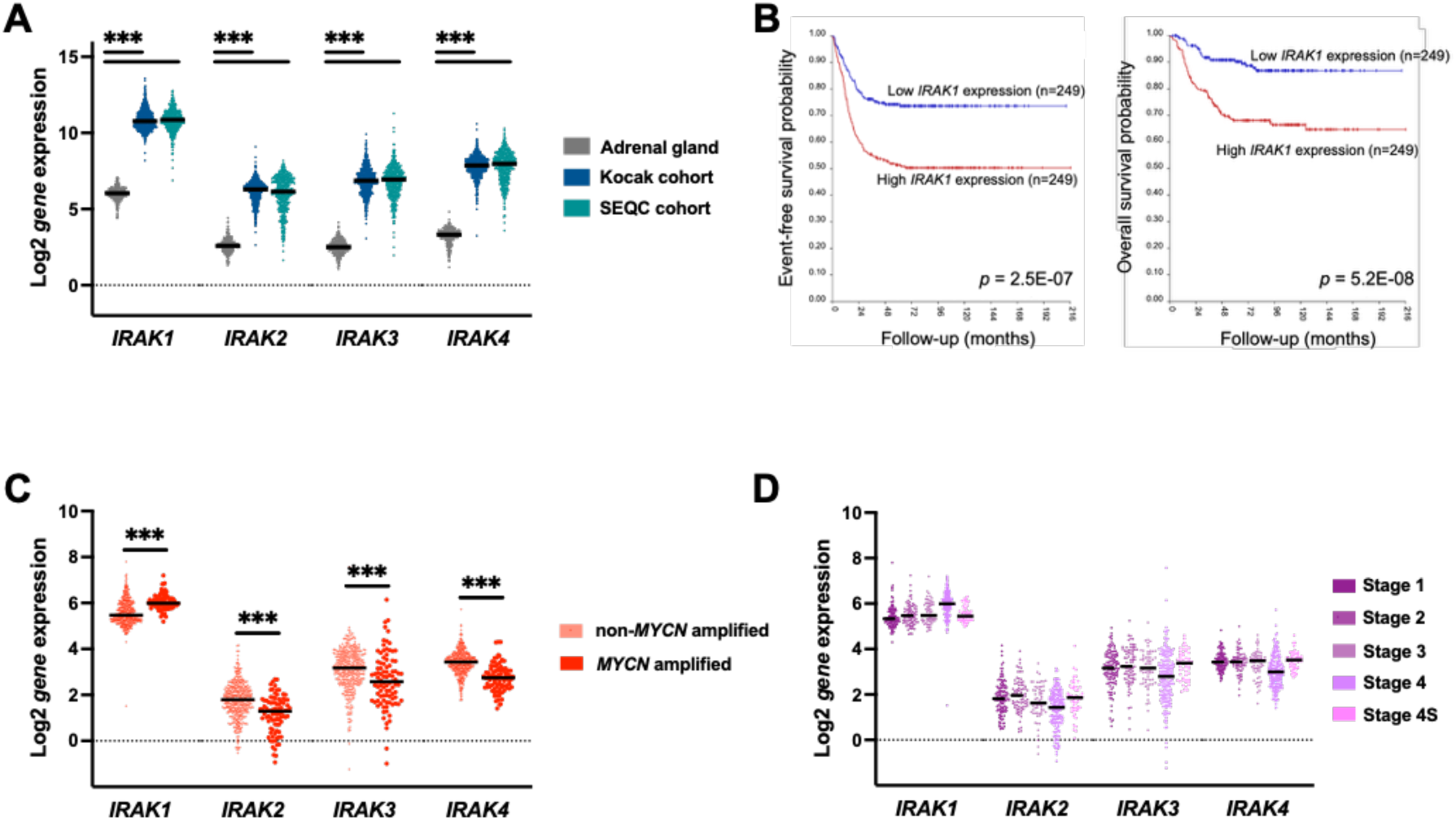
IRAK1 is a clinically-relevant target in neuroblastoma. (**A**) Normal adrenal gland tissue and NB tissue gene expression values were obtained from the GTeX databases and the SEQC and Kocak cohorts, respectively. Violin plot representation showing relative Log2-transformed gene expression for the *IRAK* isoforms in normal adrenal gland tissue (n= 295) and NB tissues (Kocak cohort (n=649) and SEQC cohort (n=498), *p*-value from two-tailed Mann–Whitney tests). (**B**) Kaplan–Meier curves showing the probability of EFS and OS for patients from the SEQC cohort. Values were dichotomized into ‘high’ and ‘low’ *IRAK1* isoform expression around the median. (**C**) *IRAK* isoform expression in NB tumor samples with (*n* = 92) and without (*n* = 406) *MYCN* amplification. The graph shows log-transformed, zero-centered expression levels obtained from microarray dataset (SEQC cohort; *p*-value from two-tailed Mann–Whitney tests). (**D**) *IRAK* isoform expression in NB tumor samples according to their diagnostic stage (stage 1, n = 121; stage 2, n = 78; stage 3, n = 63; stage 4, n = 183 and stage 4S, n =53). The graph shows log-transformed, zero-centered expression levels obtained from microarray dataset (SEQC cohort; *p*-value from two-tailed Mann–Whitney tests).

### IRAK1 regulates chemoresistance in neuroblastoma

To better understand the biological role of IRAK isoforms in NB, we first checked the gene expression of each isoform in a large panel of NB cells harboring different molecular features. This revealed that the 4 isoforms were detected in all tested NB cell lines, with *IRAK1* displaying the most consistent gene expression across the panel, while *IRAK3* expression levels varied between cell lines (Fig. 4A). Using siRNA sequences specifically targeting each isoform (Supp. 3A), our data indicated that the transient knockdown of *IRAK2* and *IRAK3* did not impact tumor spheroid viability, observed at day 8 in 3 NB cell lines (Fig. 4B). In contrast, the downregulation of *IRAK1* significantly decreased the viability of all tumor spheroid models (Fig. 4B). A significant decrease of 46 ± 15 and 27 ± 5 % was also observed following *IRAK4* downregulation in BE(2)-C and SH-SY5Y tumor spheroid viability, respectively (Fig. 4B). In addition, the knockdown of *IRAK1* and *IRAK4* expression resulted in a significant drop in the IC_50_ of vincristine from 30 ± 3 nM to 8 ± 3 nM and 8 ± 4 nM in BE(2)-C and from 38 ± 8 nM to 9 ± 1 nM and 17 ± 4 nM in SH-SY5Y following the downregulation of *IRAK1* and *IRAK4,* respectively (Fig. 4C). In the SK-N-BE(2) cell line, only the silencing of *IRAK1* expression led to a significant decrease in the IC_50_ of vincristine from 1.6 ± 0.6 nM to 0.2 ± 0.1 nM (Fig. 4C). We next performed a functional drug screen with a home-made library of 44 drugs, including 6 epidrugs, 9 repurposed drugs, 9 conventional chemotherapies and 20 targeted therapies (Supp. 3B). We performed this drug screen in three NB cell lines (BE(2)-C, SH-SY5Y, and SK-N-AS) following siRNA transfection targeting *IRAK1* (Supp. 3C-D). We used the difference in AUC between the *IRAK1* silencing and control conditions: drugs showing differences in AUC of more than 10%.mol.L^−1^ were considered to be potentiated by *IRAK1* siRNA transfection. Our data indicated that silencing *IRAK1* expression led to a chemo-sensitivity of 61, 50 and 34% of the 44-drug library in BE(2)-C, SK-N-AS and SH-SY5Y cells, respectively (Fig. 4D).

**Figure 4.**
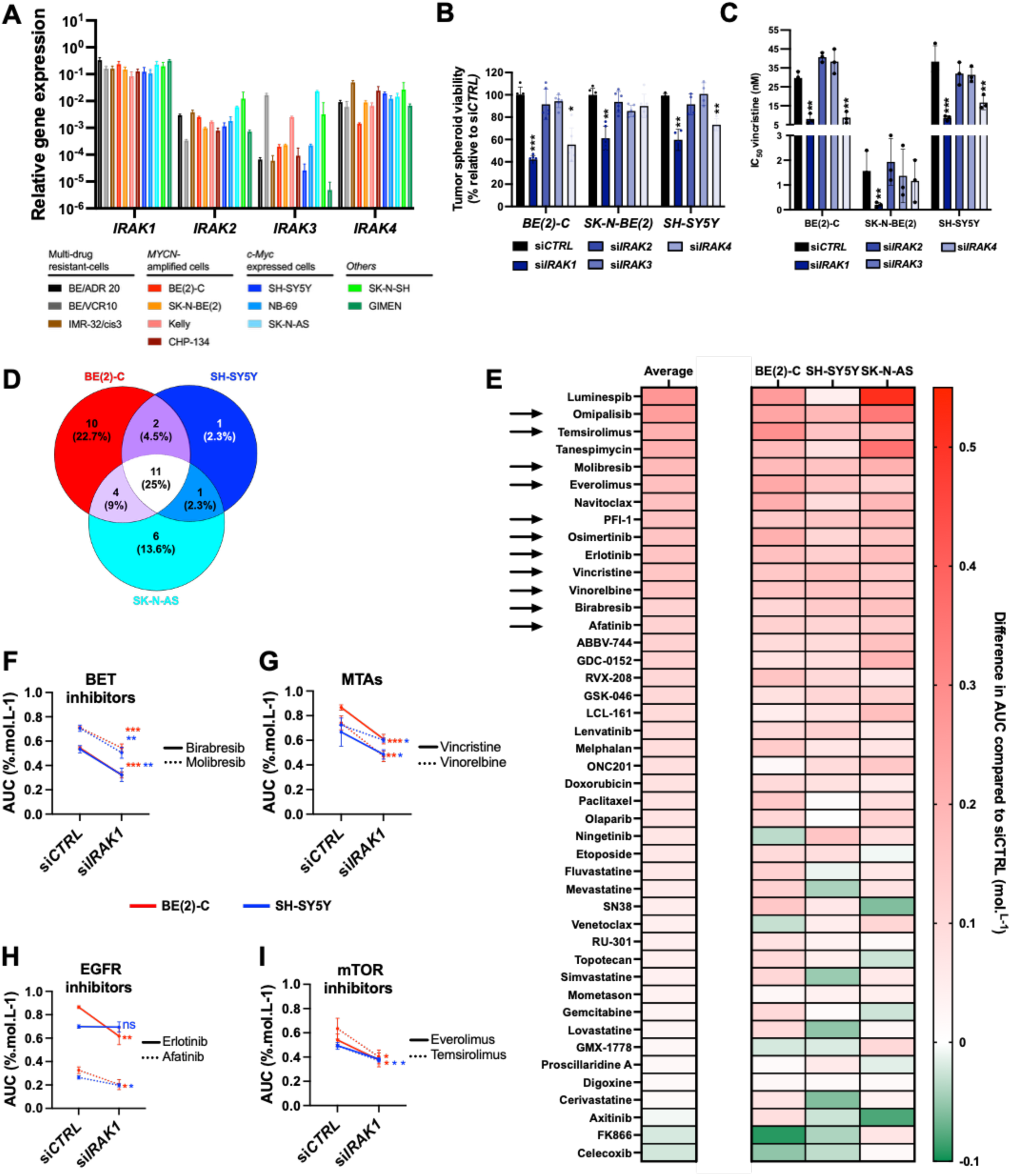
IRAK1 regulates chemoresistance in neuroblastoma. (**A**) *IRAK* isoform gene expression in a panel of 12 NB cell lines by qRT-PCR using *YWHAZ* as housekeeping gene. *Bars*, mean of at least 4 independent experiments; *Error bars*, S.D. (**B**) Histograms representing BE(2)-C, SK-N-BE(2) and SH-SY5Y tumor spheroid viability at day 8 following siRNA transfection targeting *IRAK1*, *IRAK2, IRAK3* and *IRAK4*. (**C**) Histograms representing the IC_50_ values of vincristine calculated after 7-days treatment in BE(2)-C, SK-N-BE(2) and SH-SY5Y 3D spheroid models either transfected with negative control or siRNA targeting *IRAK1, IRAK2, IRAK3* and *IRAK4*. (**D**) Venn diagram analysis of the molecules showing differences in AUC between the *IRAK1* silencing and control conditions of more than 10%.mol.L^−1^ in BE(2)-C, SK-N-AS and SH-SY5Y cell lines. (**E**) Heat map classification representing the differences in AUC between the *IRAK1* silencing and control conditions in BE(2)-C, SK-N-AS and SH-SY5Y cells lines and the average results obtained in the 3 NB cell lines for the 44 hit compounds. (**F-I**) AUC values of BETi (**F**), MTAs (**G**), mTORi (**H**) and EGFRi (**I**) in BE(2)-C and SH-SY5Y cell lines either transfected with negative control or siRNA targeting *IRAK1*. Values are the average of at least three independent experiments ± S.D; ns, *p* > 0.05; *, *p <* 0.05; *\*\*, p* ≤ 0.01 and \**\*\*, p* ≤ 0.001.

A full report of this functional screen can be found in Supplementary Table 7. A total of 11 drugs reached the 10%.mol.L^−1^ cut-off in all tested NB cell lines, including 3 BET inhibitors, 2 conventional chemotherapies and 6 targeted therapies (Fig. 4D-E). We were able to validate 8 of them in the BE(2)-C cell line, including 2 BET inhibitors (birabresib, molibresib), 2 microtubule-targeting agents (MTAs; vincristine, vinorelbine), 2 mTOR inhibitors (everolimus, temsirolimus) and 2 EGFR inhibitors (osimertinib, erlotinib; Fig. 4F-I and Supp. 3E-H). The same trend was observed in the SH-SY5Y cell line (Fig. 4F-I and Supp. 3E-H). Collectively, these results demonstrate that IRAK1 represents a chemoresistance factor that could be therapeutically exploited in NB.

### IRAK inhibition overcomes resistance to microtubule-targeting agents in neuroblastoma

To determine the therapeutic potential of inhibiting IRAK signaling in NB, we first determined the activity of the IRAK1/4 inhibitor (IRAK1/4i) in a panel of NB cell lines using a 72h dose-response assay. Our data showed that the majority of tested cell lines was relatively insensitive to IRAK1/4i at clinically-relevant concentrations, with projected IC_50_ values exceeding 100 µM in 8 out of 12 cell lines (Supp. 4A). Using another pharmacological inhibitor, HS-243 displaying a similar *in vitro* activity against IRAK1 and IRAK4 than IRAK1/4i (Supp. 4B), similar results were obtained (Supp. 4C). This suggests that the pharmacological inhibition of IRAK signaling by itself is not sufficient to reach a therapeutic effect. In order to determine whether the pharmacological inhibition of IRAK may increase the efficacy of compounds used in oncology, we performed a drug combination screen with a custom-made library of 88 FDA-approved compounds. This library comprising 13 epidrugs, 45 targeted therapies, 20 repurposed drugs and 10 conventional chemotherapies (Fig. 5A and Supplementary Table 8) was tested alone or in combination with a single dose of IRAK1/4i or HS-243 in 5 NB cell lines, harboring distinct molecular features (Fig. 5B). Drug combinations with differences in AUC of more than 10%.mol.L^−1^ were considered as potentially synergistic and below 10%.mol.L^−1^ as potentially antagonistic. Amongst the 176 pair-wise combinations tested in 5 NB cell lines, 8% appeared to have a potential synergistic effect, with the multi-resistant BE/VCR10 cell line being the most chemo-sensitized, whereas no molecule was sensitized by the IRAK inhibitors in the SK-N-AS model (Supplementary Table 8). Only 2.6% of all the drug combinations tested had a potentially antagonistic effect, confirming the role of IRAK as a chemoresistance factor in NB (Supplementary Table 8). Our results also indicated that 16 molecules showed increased efficacy when combined with both IRAK inhibitors (Fig. 5C; top right red scare). Amongst them, 7 belong to the class of microtubule-targeting agents (MTAs). In contrast, the activity of only 6 molecules was reduced when combined with both IRAK inhibitors (Fig. 5C; bottom left green scare). Regarding the 10 conventional chemotherapies tested in our library, the combination of IRAK inhibitors with most of the MTAs was highly synergistic, whereas their combination with topoisomerase inhibitors resulted in an antagonistic effect (Fig. 5D). No repurposed drug reached the 10% cut-off defining a potentially synergistic combination with the IRAK inhibitors in the tested NB models (Supp. 5D). A slight increase in the antitumor effects of the epidrug BET inhibitors when combined with IRAK inhibitors was observed (Supp. 5E), reinforcing the results of our functional genomic screen. Finally, 45 targeted therapies were tested covering major signaling pathways involved in cancer. Our drug combination screen did not highlight a specific signaling pathway that could synergize with the inhibition of IRAK signaling in all NB cell lines (Supp. 5F).

**Figure 5.**
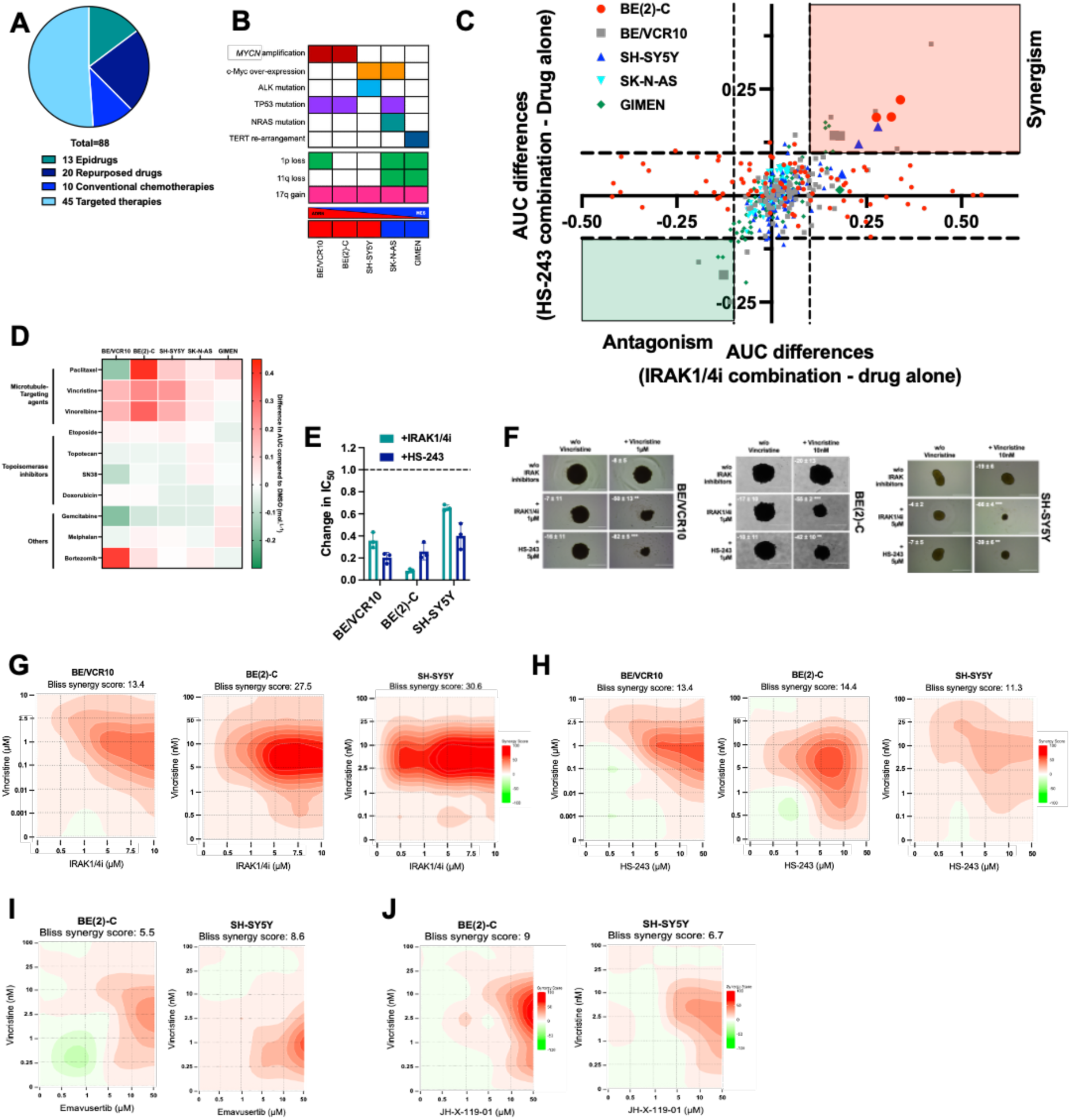
Inhibition of IRAK overcomes resistance to microtubule-targeting agents in neuroblastoma. (**A**) Eighty-eight compounds were used for the drug combination screening and are represented in donut diagram and classified by pharmacological classes. (**B**) Molecular and genetic features of the 5 NB cell lines used for the drug combination screening. (**C**) Differences in AUC between monotherapy and combination with IRAK inhibitors, IRAK1/4i (x axis) or HS-243 (y axis) for all 88 compounds of the drug library and in the 5 NB cell lines used after 72h drug incubation. (**D**) Heat map classification representing the differences in AUC between the average of both IRAK inhibitors and DMSO conditions in BE/VCR10, BE(2)-C, SH-SY5Y, SK-N-AS and GIMEN cells lines for the 10 conventional chemotherapy agents. (**E**) Histogram representation of changes in IC_50_ values in 3 NB spheroid models when vincristine is used in combination with subtoxic concentrations of IRAK inhibitors at day 7. (**F**) Representative photographs of BE/VCR10 (left panel), BE(2)-C (middle panel) and SH-SY5Y (right panel) tumor spheroids treated with vincristine or IRAK inhibitors alone and the combination of both at day 7. Mean decrease in spheroid viability is indicated ± S.D. Scale bar, 1 mm. All values are the average of at least three independent experiments ± S.D. (**G, H**) A 6 × 5 matrix was used to test drug combinations between vincristine and (**G**) IRAK1/4i or (**H**) HS-243 in BE/VCR10 (left panel), BE(2)-C (middle panel) and SH-SY5Y (right panel) tumor spheroid models. Heat maps representing the Bliss score obtained at day 7. (**I, J**) A 6 × 5 matrix was used to test drug combinations between vincristine and (**I**) Emavusertib or (**J**) JH-X-119-01 in BE(2)-C (left panel) and SH-SY5Y (right panel) tumor spheroid models. Heat maps representing the Bliss score obtained at day 7.

Given that the combination of MTAs with IRAK inhibitors represents the most promising therapeutic strategy, we focused our validation on vincristine, a drug used in the clinic to treat NB patients. To validate the results from the drug combination screen in 3D spheroid models, a minimally cytotoxic concentration of each IRAK inhibitor was combined with a range of concentrations of vincristine. Our data showed that both IRAK inhibitors sensitized the BE/VCR10, BE(2)-C and SH-SY5Y cells to vincristine, as evidenced by a significant decrease in its IC_50_ (Fig. 5E-F). Indeed, a small decrease in the BE/VCR10 spheroid viability was observed with monotherapies of 1 µM vincristine, 1 µM IRAK1/4i and 5 μM HS-243 (8.3 ± 5.2%, 6.7 ± 10.7% and 16.1 ± 11.1%, respectively, Fig. 5E-F), whereas the combination with IRAK1/4i and HS-243 led to a significant decrease in BE/VCR10 spheroid viability compared to untreated cells (50.3 ± 13.4% and 81.8 ± 4.6%, respectively, Fig. 5E-F, p < 0.01). Similar results were obtained in the parental BE(2)-C cell line and the SH-SY5Y cell line (Fig. 5E-F). Next, we performed a 6 × 5 matrix that contained the two compounds at different concentration ratios, allowing to cover a large range of drug combinations. To assess the synergy of the two compound treatments, the Bliss score was calculated^25^. As expected from both our pharmacological and functional screen results, the combination of vincristine and IRAK1/4i produced a strong synergistic effect in all 3D spheroid models tested with an overall Bliss score of 13.4, 27.5 and 30.6 in BE/VR10, BE(2)-C and SH-SY5Y cells, respectively (Fig. 5G). Similar results were obtained with the HS-243 inhibitor (Fig. 5H). The development of IRAK inhibitors in clinical oncology is currently emerging with dual inhibitor targeting IRAK1 and IRAK4 but also, selective inhibitors of each isoform ^22^. In two NB spheroid models, selective IRAK1 or 4 inhibitors (JH-X-199-01 and Emavusertib, respectively) showed reduced synergism with vincristine as compared to dual IRAK1/4 inhibitors (Fig. 5I-J), suggesting that dual inhibition of both IRAK 1 and 4 is necessary to increase the chemo-sensitivity of vincristine in NB.

### IRAK1/4 inhibition synergizes with vincristine in *ex vivo* and *in vivo* neuroblastoma models

To assess the efficacy of a therapeutic strategy based on IRAK inhibition in preclinical models that can accurately predict outcomes in the clinic ^26^, we first tested the combination of IRAK1/4i with vincristine in established 3D NB tumoroids as previously described ^27^. As expected, our data showed a dose-response effect of vincristine in all tested models while no response was observed with the IRAK1/4i (Supp. 5A-B). Nevertheless, combining IRAK1/4i with vincristine resulted in a significant increase in cell death in all tested NB organoids. Indeed, the drug combination led to a significant increase from 28.3 ± 5.1 % in cell death in the PDX NB1 treated with vincristine alone to 42.3 ± 10.3 % when combined with 1 µM of IRAK 1/4i (Fig. 6A, p < 0.001). Using a fluorescent live / dead staining assay, the results were confirmed in 2 NB organoid models with a significant increase of dead cells following the drug combination treatment (Fig. 6B-C, p < 0.01). Finaly, in order to mimic tumor relapse following initial response, we performed a cell regrowth experiment (Fig. 6D). While vincristine as monotherapy is insufficient itself to block the regrowth of NB organoid models, its combination with IRAK1/4i is highly effective to strike the regrowth down in the PDX-NB2 and NB3 models (Fig. 6E-G). Then, to increase the complexity of our preclinical model and strengthen our results, we tested the drug combination in a syngenic orthotopic mouse model, using a *MYCN*-driven NB cell line derived from TH-MYCN transgenic mice. Despite the lack of anti-tumor activity of the IRAK1/4i itself, its combination with a low dose of vincristine led to a significant decrease in tumor growth in comparison to vehicle (Fig. 6H). This was reflected by a reduction in the tumoral mass measured by micro ultra-sounds (Fig. 6I). Finally, as shown in Fig. 6J, the median survival of mice treated with single agents was not different in comparison to the vehicle-treated mice, whereas the drug combination resulted in a significant increase in median survival (28 *versus* 22 days, p < 0.05). Altogether, our study pinpoints the clinical relevance of using IRAK1/4 inhibitors in combination with conventional chemotherapy in NB.

**Figure 6.**
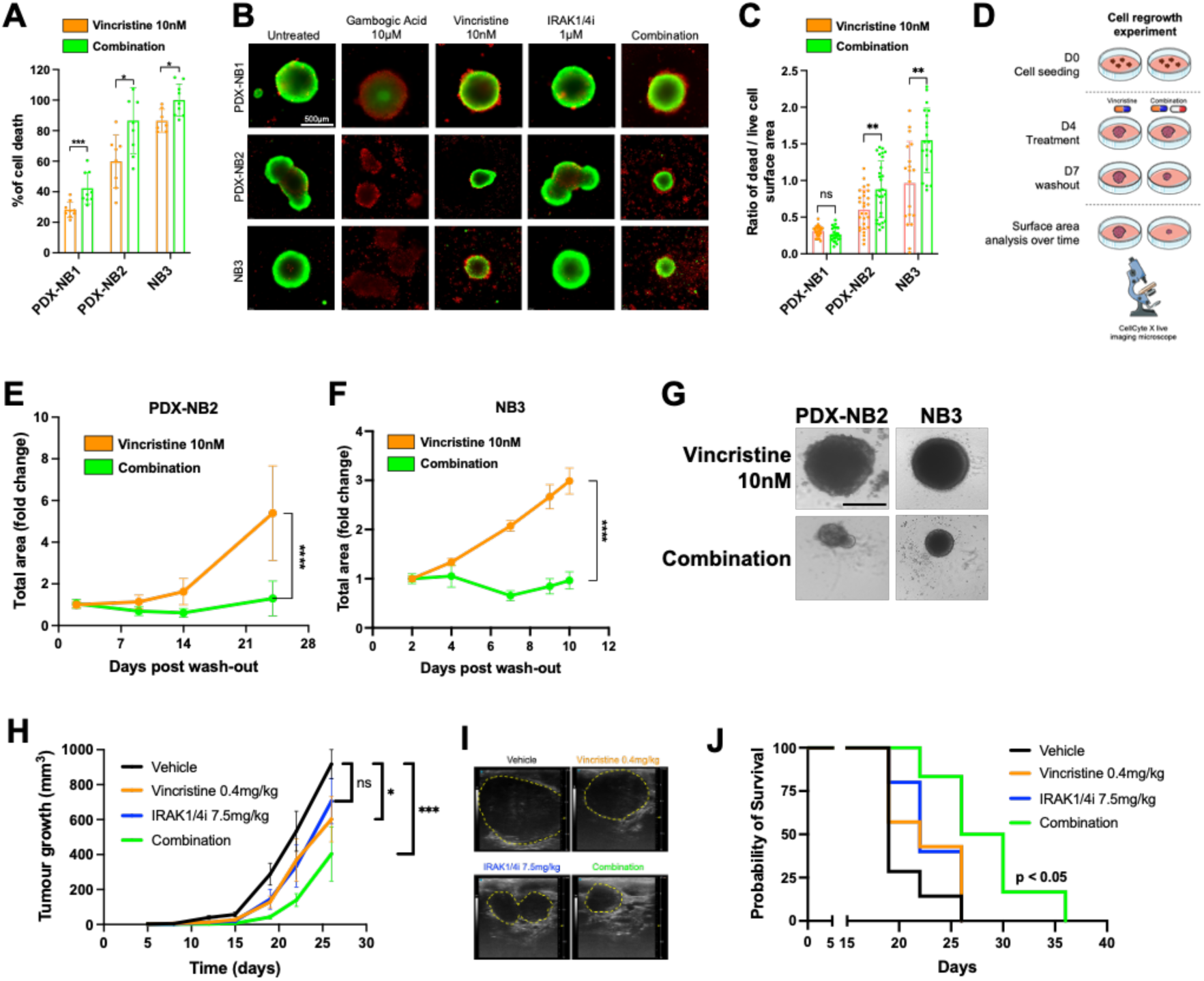
IRAK inhibitor synergizes with vincristine in *ex vivo* and *in vivo* neuroblastoma models. (**A**) Cell death (%) comparison across NB organoids treated with vincristine (10nM) alone or in combination with IRAK1/4i (1μM) at 72h. Cell death is expressed as percentage relative to CellTox-Green-treated control (TritonX100; 0.005%). Values represent the mean of at least three independent experiments ± S.D. *, p < 0.05; ***, p ≤ 0.001 (Mann-Whitney test). (**B**) Representative LIVE/DEAD staining (calcein-AM/propidium iodide) of NB organoids 72h post-treatment with vincristine (10nM), IRAK1/4 inhibitor (1μM), or combination therapy. Green fluorescence indicates viable cells; red fluorescence indicates dead cells. Scale bar: 500 μM. (**C**) Quantitative analysis was performed by measuring the ratio between viable (green) and dead (red) cell surface areas using ImageJ/Fiji software. Values represent the mean of at least 20 independent organoids from two (NB3) to three (PDX-NB1; PDX-NB2) independent experiments ± S.D. ns, p<0.5; **, p ≤ 0.01. (**D**) Workflow of the cell regrowth experiment. (**E, F**) Time-course analysis of NB organoid growth following drug treatment with vincristine (10nM) alone or combined with IRAK1/4i (1μM) and subsequent washout. Total NB organoid area was measured over 10-24 days post-treatment in PDX-NB2 (**E**) and NB3 (**F**) organoids. Experimental endpoints were selected to allow control NB organoids to reach optimal size for expansion (800-1,000 μM). Follow-up monitoring and curve generation (area fold change compared to day 0 (D0) post-washout) were automated using a live cell imaging system. Values represent the mean of at least 10 independent organoids from one (NB3) to two (PDX-NB2) independent experiments ± S.D. ****, p < 0.0001 (Mann-Whitney test). (**G**) Representative brightfield images show NB organoids at the final timepoint for both tested conditions. Scale bar: 500 μM. (**H**) Eight-week-old C57/BL6 mice were orthotopically injected with 0.5 million 9464D cells, a *MYCN*-driven NB cell line derived from TH-MYCN transgenic mice into the left adrenal gland. All treatments were administered by intraperitoneal injection every 3 days until endpoint. Tumor volumes were measured twice a week using micro ultra-sounds. Statistical significance was tested using unpaired Student’s t test. Significant differences compared to vehicle. (**I**) Representative pictures of tumors taken by ultra-sounds and processed in an *in vivo* analysis software Vevo LAB. (**J**) Kaplan-Meier survival curves (n ≥ 5/group) from the different groups of treatment. The log-rank test was used to compare survival rates by univariate analysis.

### IRAK inhibition enhances the vincristine anti-cancer properties independently of the IRAK canonical axis

To better understand the mechanism of action of the drug combination, we conducted single cell RNA-seq (scRNA-seq) on NB cells pre- and post-treated with vincristine alone, IRAK1/4i alone or the combination of both using the 10x Chromium platform, performing analyses individually and in an integrated manner. Differential gene expression analysis showed that in comparison to DMSO control sample, all the treated conditions are very similar at the transcriptomic level, as reflected by a high number of shared differentially expressed genes across the conditions (Fig. 7A). These data indicated that there is no treatment-induced cell type-specific molecular dynamics. Next, our analysis identified 4 different clusters shared with all conditions, while a fifth cluster appeared only in the high dose vincristine-treated sample as well as drug combination samples (Fig. 7B). However, no specific signaling pathway emerged from this cluster 5 compared to the other one by performing GSEA (Supp. 6A). Focusing on the cluster 5 comparison between high dose vincristine-treated sample and the combination of low dose vincristine with IRAK1/4i at 5µM, although our data revealed 118 differentially-expressed genes (Supp. Table 10), no significant dysregulated pathway was identified. Altogether, these results point out that the drug combination induced overall similar transcriptomic changes than vincristine high dose.

**Figure 7.**
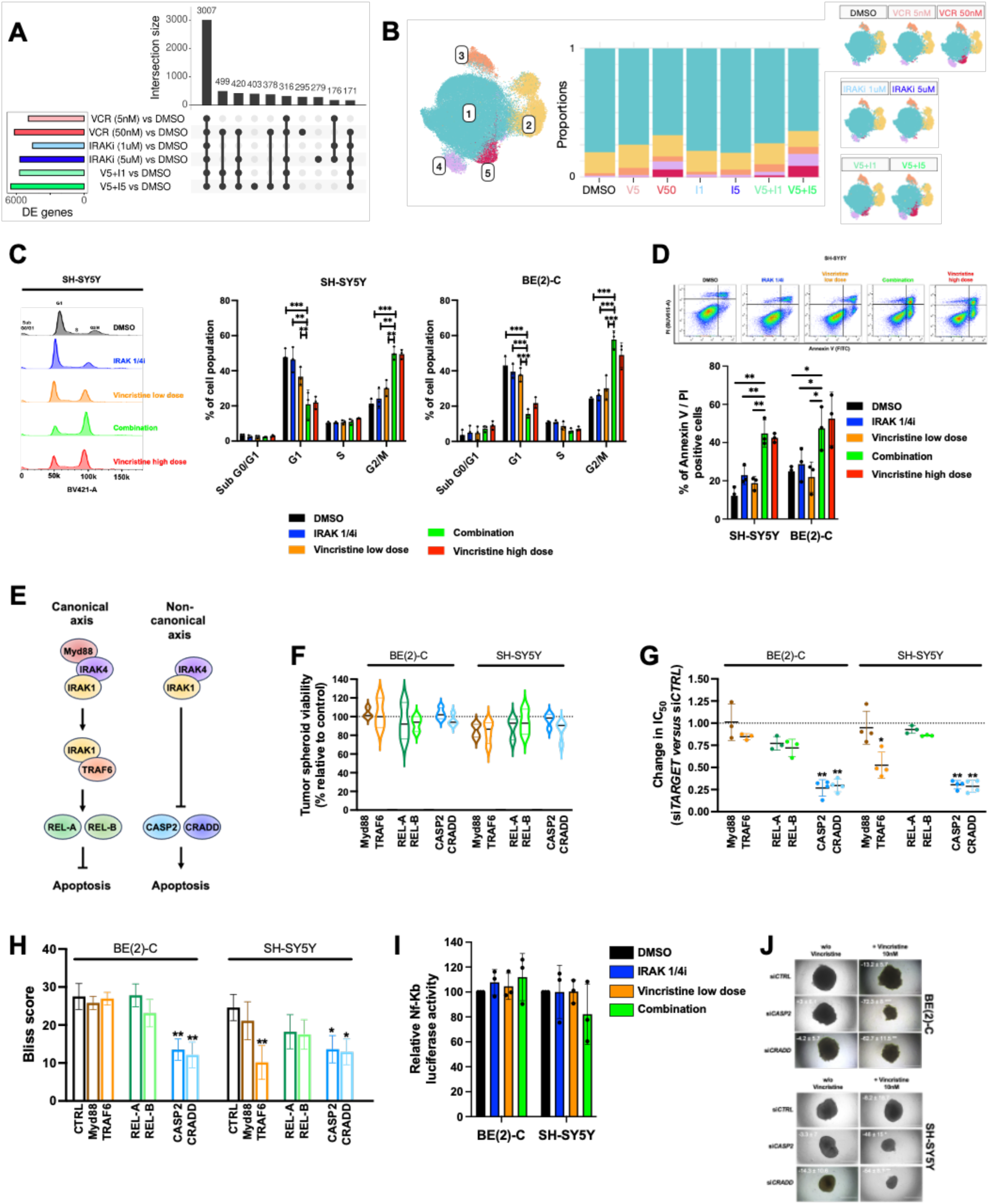
IRAK inhibition enhances the pro-apoptotic activity and cell cycle arrest of vincristine independently of the IRAK canonical immune pathway. **(A)** Upset plot presenting overlaps of differentially expressed genes for each condition vs control (DMSO). The barplot on top indicates the size of each gene set intersection, with the included conditions displayed in the matrix below (only intersections of 150+ genes are shown). The barplot on the left shows the total number of DE genes per comparison. (**B**) Overview of cell clusters and their distribution across samples. Included are the UMAP embedding of all cells colored by cluster identity, split by sample below. The barplot represents the proportion of cells per cluster in each sample. (**C**) Representative cell cycle profiles of SH-SY5Y cells incubated for 8 h with IRAK1/4i (1 µM – blue), vincristine low or high dose (10 nM or 50 nM – orange or red) or the combination of IRAK1/4i and vincristine low dose (green). Histograms representing the cell cycle phase distribution in SH-SY5Y and BE(2)-C with the indicated treatments. (**D**) Representative dot plots of SH-SY5Y cells following Annexin V-FITC (x axis) and PI (y axis) staining after 24-h incubation with IRAK1/4i (1 µM – blue), vincristine low or high dose alone (10 nM or 50 nM – orange or red) or the combination of IRAK1/4i and vincristine low dose (green). Histograms showing the annexin-V / PI positive cells after vincristine and/or IRAK1/4i treatment in SH-SY5Y and BE(2)-C cell lines. (**E**) Representation of the canonical and non-canonical axis of IRAK1. (**F**) Box plot representing BE(2)-C and SH-SY5Y tumor spheroid viability at day 8 following siRNA transfection targeting *Myd88*, *TRAF6, REL-A, REL-B, CASP2* and *CRADD*. (**G**) Histogram representation of changes in IC_50_ values in BE(2)-C and SH-SY5Y tumor spheroid models when vincristine is used in combination with siRNA targeting *Myd88*, *TRAF6, REL-A, REL-B, CASP2* and *CRADD* at day 7. (**H**) Histogram representation of the Bliss score calculating the synergy between vincristine and IRAK1/4i at day 7 in BE(2)-C and SH-SY5Y tumor spheroid models transfected with siRNA targeting *Myd88*, *TRAF6, REL-A, REL-B, CASP2* and *CRADD*. (**I**) Histogram representation of the NF-KB activity in BE(2)-C and SH-SY5Y cells following treatment with vincristine alone, IRAK1/4i alone and their combination. (**J**) Representative photographs of BE(2)-C (top panel) and SH-SY5Y (bottom panel) tumor spheroids treated with vincristine and the combination of siRNA targeting *CASP2* and *CRADD* at day 7. Mean decrease in spheroid viability is indicated ± S.D. Scale bar, 1 mm. All values are the average of at least three independent experiments ± S.D. Values are the average of at least three independent experiments ± S.D; ns, *, *p <* 0.05; *\*\*, p* ≤ 0.01 and \**\*\*, p* ≤ 0.001.

Also, given that MTAs are known to inhibit cancer progression by triggering cell cycle arrest and programmed cell death, we first analyzed whether IRAK1/4i could potentiate vincristine efficacy through these mechanisms. Cell cycle analysis revealed that IRAK1/4i increased the mitotic arrest induced by vincristine in SH-SY5Y and BE(2)-C cells after 8h and 24h drug incubation, respectively (Fig. 7C). This is reflected by an increase in the percentage of cells arrested in G2/M from 30.1 ± 4.6 to 49.7 ± 3.8 % and from 30.1 ± 7 to 57.7 ± 5.2 % in SH-SY5Y and BE(2)-C cell lines, respectively treated with vincristine alone or its combination with IRAK1/4i (Fig. 7C; *p* < 0.001). Consistently, treatment with 1 µM IRAK1/4i alone did not induce apoptosis in NB cell lines but it increased the level of Annexin V / PI positive cells induced by 10 nM vincristine from 18.8 ± 3.5 to 44.8 ± 7.1 % and from 22 ± 7.9 to 47.5 ± 11.4 % when used in combination in SH-SY5Y and BE(2)-C cells, respectively (Fig. 7D; *p* < 0.05). Then, we studied the role of the canonical IRAK signaling pathway in the mechanism of action of the drug combination (Fig. 7E). Using transient knockdown of the adaptor protein MyD88, which bridges IRAK1/4 to IL-1R/TLRs and is essential for IRAK1/4 activation in innate immunity ^24^, the proximal effector TRAF6 as well as the downstream targets REL-A and REL-B (Supp. 6B), we did not observe any impact on tumor spheroid viability, vincristine sensitivity as well as drug combination synergism, with the exception of the downregulation of *TRAF6* in the SH-SY5Y cell line leading to a significant decrease in vincristine sensitivity and a drop of the drug combination Bliss score (Fig. 7F-H). Moreover, NF-κB activity was assessed following the different treatment conditions in BE(2)-C and SH-SY5Y cell lines. Our data showed no significant dysregulation of NF-κB activity in all tested conditions (Fig. 7I). Since previous studies demonstrated a non-canonical activity of IRAK1 through the PIDDosome complex ^20, 28^, we tested the impact of *CASP2* and *CRADD* gene silencing (Fig. 7E). While no significant effect was observed on tumor spheroid viability (Fig. 7F), an increased sensitivity to vincristine and a partial rescue of the combinatorial effect was shown in both models (Fig. 7G-H, J). Collectively, our results demonstrated that IRAK inhibition can increase vincristine antitumor activity through its non-canonical axis involving the PIDDosome complex.

## DISCUSSION

High-risk NB remains one of the most challenging pediatric cancers, where relapse and chemoresistance lead to dismal survival rates. Leveraging pharmacological and functional high-throughput screening, chemo-informatics and transcriptomics analyses, our study reveals that IRAK1 represents a major chemoresistance factor in NB, and provides a strong foundation for future clinical trials targeting IRAK signaling in high-risk NB patients.

Multiple large-scale programs have been developed across the globe, such as Zero Childhood Cancer (ZERO), INdividualized Therapy FOr Relapsed Malignancies in Childhood (INFORM) or Molecular Profiling for Pediatric and Young Adult Cancer Treatment Stratification (MAPPYACTS) in order to unveil functional vulnerabilities in pediatric cancers, including NB ^29–31^. Although these programs have demonstrated that comprehensive and entity-agnostic pediatric precision oncology is feasible, they also highlighted the limitations of personalized treatments solely based on molecular tumor profiling. Indeed, genomic analysis of NB tumors has indicated a low rate of mutations in druggable targets; with the exception of anaplastic lymphoma receptor tyrosine kinase (ALK), there are no kinases mutated at a rate greater than 5% ^32,33^. Using high-throughput screening, several studies strove for the identification of non-mutated targetable vulnerabilities in NB such as microRNA or epigenetic-driven changes ^34–36^ or for the development of drug combination strategies ^37–39^. In this study, we aimed at uncovering chemoresistance factors in NB by establishing a reverse molecular pharmacology approach and developing biology-guided combinatorial treatments to overcome drug resistance. Based on the concept of drug poly-pharmacology using commercially available libraries of compounds, we hypothesized that the annotated targets of an identified chemosensitizer hit may be involved in modulating resistance, and could thus represent a targetable vulnerability. Our results indicated that gefinitib, a first-class EGFR inhibitor was able to increase the efficacy of vincristine in a multi-resistant NB cell line. Performing a target deconvolution strategy using chemo-informatics databases, we found that gefinitib has 14 known targets, including IRAK1 and IRAK4.

The IRAK family plays a crucial role in innate immunity by participating in interleukin-1 and TLR-mediated signaling pathways. It is composed of IRAK-1, -2, and -4, which are expressed in a variety of human cell types and IRAK-M (−3) whose expression is largely limited to monocytes and macrophages ^40^. Over the last decade, cancer-specific dependencies on IRAK signaling have started to emerge. Past studies suggest that elevated IRAK1 or IRAK4 expression levels correlate with more aggressive disease, including hepatocellular carcinoma, pancreatic and lung cancers as well as hematological malignancies ^19,41–43^. Here, we observed a significant upregulation of all IRAK members in NB tissues. However, only the high expression of *IRAK1* correlates with poor clinical outcome, independently of established prognostic indicators for NB. Moreover, the current study not only identifies *IRAK1* expression as a powerful independent prognostic indicator of poor clinical outcome in NB patients, but also shows that IRAK1 plays a key role as a chemoresistance factor, which could explain its prognostic value. Taking advantage of a functional drug screening, we demonstrated that downregulation of IRAK1 expression increased the efficacy of drugs from several therapeutic classes, including epidrugs such as BET inhibitors, targeted therapies such as mTOR, EGFR or HSP90 inhibitors as well as conventional chemotherapies. These results are consistent with previous studies demonstrating the key role of IRAK1 expression in chemoresistance in breast, endometrial and hepatocellular carcinoma ^17,19,44^. In addition, its role in radioresistance has also been described in breast and cervical cancers as well as gliomas ^20,28,45^.

Among the 4 IRAK family members, IRAK1 and IRAK4 are the only ones to exhibit serine/threonine kinase activity and are therefore targetable by small-molecule inhibitors ^24^. Since monotherapies are highly unlikely to cure patients, to quickly reveal potentially synergistic drug combinations with IRAK inhibitors, we performed a drug combination screen in five NB cell lines using two dual IRAK1/4 inhibitors. Our previous study and others demonstrated that drug synergism is rare, occurring in less than 5% of all tested combinations ^10,46,47^. Here, amongst the 176 tested pair-wise combinations, 8.4% appeared to have a potentially synergistic effect, with a rate reaching 23.9% in the multi-resistant NB cell line, confirming the major role of IRAK1 as a chemoresistance factor. Using high-throughput screening, our study underlined a highly synergistic effect between microtubule-targeting agents and the functional or pharmacological inhibition of IRAK1 in NB. These results were confirmed in clinically-relevant NB models including 3D spheroids, patient-derived tumoroids and an orthotopic syngenic mouse model. Previous studies also highlighted the therapeutic potential of combining IRAK inhibition with microtubule-targeted agents to overcome drug resistance in breast and nasopharyngeal cancers as well as in leukemia ^17,21,48^. Originally developed for the treatment of chronic inflammatory auto-immune diseases such as rheumatoid arthritis ^16^, only two IRAK inhibitors are currently tested in oncology. CA-4948, also called Emavusertib, a multi-kinase inhibitor of IRAK4 and FLT3 is currently being evaluated in a phase 1/2 clinical trial in acute myeloid leukemia and myelodysplastic syndrome as a monotherapy [NCT04278768] and in combination with conventional chemotherapies and immunotherapies in advanced or metastatic colorectal cancers, biliary tract carcinoma and urothelial cancers [NCT06696768; NCT07107750; NCT06439836]. The second IRAK inhibitor, R289, is a dual IRAK1/4 inhibitor currently tested as a monotherapy in a phase 1b in relapsed / refractory lower risk myelodysplastic syndrome. To date, few selective IRAK1 inhibitors are in preclinical development, including the covalent IRAK1 inhibitor, JH-X-199-01, but none of them are under clinical investigation in oncology ^49^. In our study, we tested selective IRAK1 or 4 inhibitors as well as dual inhibitors. Our results indicated that the dual inhibition of IRAK1 and 4 led to a stronger synergistic combination with the microtubule-targeting agents.

The MyDDosome complex is required for the activation of IRAK1/4 in innate immunity leading to the activation of its proximal effector TRAF6 and then the NF-KB pathway ^24,40^. Our data describe a mechanism through which IRAK1/4 inhibition increased NB cell chemosensitivity in a MyD88-independent manner. Recent studies revealed a non-canonical pathway of IRAK signaling as playing a key role in response to irradiation ^20,50^. Rather than involving the TLR/Myd88/TRAF6 axis, in this context IRAK1 interacts with the pro-apoptotic PIDDosome complex, leading to cell survival. In our study, we found that downregulation of the expression of key members of the PIDDosome complex, *RAIDD* and *CASP2*, sensitized NB cells to vincristine and decreased the efficacy of the drug combination, while *Myd88* or *REL-A/B* silencing did not have any impact. These results reveal a new role of the non-canonical IRAK1 pathway in NB chemosensitivity.

Altogether, our study uncovers the role of IRAK1 as a chemoresistance factor in NB and its dual inhibition with IRAK4 combined with conventional chemotherapies represents a potential therapeutic avenue in NB, providing a strong foundation for future clinical trials. Our findings also constitute a proof-of-concept of our reverse molecular pharmacology approach to quickly identify chemoresistance factor and develop biology-guided drug combinations for refractory cancers.

## MATERIALS AND METHODS

### Cell culture

SK-N-AS (RRID:CVCL_1700), SH-SY5Y (RRID:CVCL_0019), SK-N-SH (RRID:CVCL_0531), NB-69 (RRID:CVCL_1448) and Kelly (RRID:CVCL_2092) NB cell lines were obtained from European Collection of Cell Cultures. SK-N-BE(2) (RRID:CVCL_0528), SK-N-BE(2)-C (referred as BE(2)-C, RRID:CVCL_0529), GIMEN (RRID:CVCL_1232) and CHP-134 (RRID:CVCL_1124) cell lines were provided by the Children’s Cancer Institute, (Sydney, Australia). BE/VCR10 and BE/ADR20 (vincristine and doxorubicin highly resistant cells derived from BE(2)-C respectively) and IMR-32/Cis3 (cisplatin highly resistant cells derived from IMR-32 (RRID:CVCL_0346)) resistant NB cell lines were kindly given by Prof. M. Kavallaris (Children’s Cancer Institute, Sydney, Australia) and Dr C. Flemming (Children’s Cancer Institute, Sydney, Australia). Upon receipt, cell master stocks were prepared and cells for experiments were passaged for less than 3 months. NB cell lines were cultured in DMEM or RPMI-1640 media (Life Technologies) supplemented with 10% FBS (#F0392; Sigma-Aldrich, GE), 1% sodium pyruvate (#11360070, Thermo-Fisher) and 1% penicillin/streptomycin (#15070063, Thermo-Fisher). Cells were cultured at 37°C and 5 % CO_2._ All cell cultures were monthly screened to ensure the absence of mycoplasma contamination (Eurofins Genomics or kit MycoAlert® Assay LONZA #LT07-710 and #LT07-518).

### Drugs and Reagents

Stock solutions of vincristine (MedChemExpress HY-N0488, USA), vinorelbine (MedChemExpress HY-12053A, USA), everolimus (MedChemExpress HY-10218, USA), temsirolimus (MedChemExpress HY-50910, USA), erlotinib (MedChemExpress HY-50896, USA), afatinib (MedChemExpress HY-10261USA), birabresib MedChemExpress HY-15743, USA), molibresib (MedChemExpress HY-13032B, USA), IRAK1/4 inhibitor (#s6598, Selleckchem, USA), HS-243 (#s9701, Selleckchem, USA), Emavusertib (MedChemExpress HY-135317, USA) and JH-X-199-01 (MedChemExpress HY-103017, USA) were prepared in dimethyl sulfoxide (DMSO). Stock solutions were kept at −20 °C and were freshly diluted in the culture medium for experiments.

### High-throughput drug screening

Three libraries (Prestwick chemical, Lopac and Tocris) containing ∼2,800 unique FDA-approved drugs and pharmacologically-active molecules (Supp. Table 1) were screened on BE/VCR10 cells at a single dose of 5µM alone or in combination with 1µM of vincristine. Cell viability was assessed after 72h incubation using a home-made Alamar Blue solution (75mg resazurin, 12.5mg methylene blue, 164.5mg potassium hexacyanoferrate III and 211mg potassium hexacyanoferrate II trihydrate, 500ml water; Sigma) as previously described ^51^. Assay performance was evaluated by calculating the Z-factor for each assay plate, according to the method described by Zhang *et al* ^52^. Compounds were considered as cytotoxic drug candidates when an inhibition of cell viability of more than 75 % in comparison to DMSO is observed and as potential chemosensitizers to overcome the resistance to vincristine when they re-sensitize BE/VCR10 cells to vincristine by at least 25%. A secondary drug combination screen using the same methodology was performed with the 91 chemosensitizers identified. Compounds were defined as confirmed hits when they re-sensitize BE/VCR10 cells to vincristine by at least 25% in both screens.

### Functional validation assay

The NB cells were seeded in T25 cell culture flasks and transfected with 1mL of Opti-MEM medium (Gibo, #11058021) containing 1% of lipofectamine RNAiMax (Life Technologies, #13778150) and 5nM of siRNA. A pool of three different siRNA sequences targeting *IRAK1*, *RET, STK33, PRMT1, PGRMC1, PPP1CA, EBP, IRAK2*, *IRAK3*, *IRAK4*, *Myd88*, *TRAF6*, *CASP2* and *CRADD* were used as well as a non-targeting negative control siRNA with no significant sequence homology to mouse, rat or human gene sequences (Silencer Select; Thermo-Fisher Scientific; Supp. Table 9). Two days later, cells were seeded in 384-well flat or U bottom and low-binding plates (#3764 or #3830, Corning, USA) in DMEM. Cell and spheroid viability were then evaluated after 3 or 8 days, respectively using Cell Titer-Glo^®^ 2.0 Cell Viability Assay (Promega, #G9243). Measurements were performed with a PHERAstar^®^ plate reader (RRID:SCR_027001; BMG). To evaluate the level of gene knock-down, cells were harvested 48h after transfection and total RNA was extracted using RNeasy Plus mini kit (Qiagen, #74136), following the manufacturer’s instructions. Reverse transcription was performed with ReadyScript^®^ cDNA synthesis kit (Sigma-Aldrich, #RDRT-100RXN) and qRT-PCR was undertaken using SsoAdvanced Universal SYBR^®^ Green Supermix (Bio-Rad) and a CFX384 Real-Time System device (RRID:SCR_018057; Bio-Rad). Gene expression levels were determined using the ΔCt method, normalized to the *YWHAZ* or *GAPDH* control genes. Predesigned SYBR^®^ Green primers (Merck, Fontenay-sous-bois, France) were used (Supp. Table 9). To evaluate the level of protein knockdown, cells were lysed in RIPA buffer (Thermo-Fisher, #89901) freshly supplemented with a cocktail of proteases and phosphatases inhibitors (Thermo-Fisher, #78442), 72h after siRNA transfection. Protein concentrations were determined using the Bio-Rad Protein Assay (Bio-Rad, #5000001). Proteins were separated by SDS-PAGE and electro-transferred onto a nitrocellulose membrane. Primary antibodies used were directed against GAPDH (clone 6C5; 1/50,000; #ab8245; Abcam; RRID:AB_2107448) and IRAK1 (1/1,000; #4359S; Cell Signaling; RRID:AB_490853). Peroxydase-conjugated secondary antibodies (Cell Signaling, #7074; RRID: AB_2099233 and #7076; RRID: AB_330924) and chemiluminescence detection kit (Millipore) were used for visualization with ChemiDoc Touch Imaging System (RRID:SCR_019037; BioRad). Membranes were probed again with GAPDH antibody following incubation with a stripping solution (#21059, Thermo-Fisher).

### Drug combination and functional screens

Briefly, 4,000 living cells were seeded per well with the Certus Flex^®^ (GyGer) in 384-well plates (Corning, #3764) for the drug combination screen. Following two-days transfection with siRNA targeting *IRAK1*, 4,000 living cells from the BE(2)-C, SH-SY5Y and SK-N-AS cell lines were seeded per well with the Certus Flex^®^ (GyGer) in 384-well plates (Corning, #3764) for the functional screen. Cells were incubated in the presence of a custom-made drug library containing 88 drugs alone or in association with a single dose of IRAK1/4i or HS-243 for the drug combination screen and in the presence of a custom-made drug library containing 44 drugs for the functional screen (Supp. Tables 7 and 8). Drugs were distributed with an Echo 550 liquid dispenser^®^ (RRID:SCR_027476; Labcyte) at 6 different concentrations covering 3 logs in constant DMSO with 3 different ranges depending on the cytotoxicity of the compound (*i.e.*, 1nM to 1μM, 10nM to 10μM or 100nM to 100μM). Cell viability was measured using CellTiter-Glo^®^ 2.0 Cell Viability Assay (Promega, #G9243) after 72h of drug incubation in a humidified environment at 37°C and 5% CO_2_. Luminescence was measured using a PHERAstar^®^ plate reader (RRID:SCR_027001; BMG). Data were normalized to negative control wells (DMSO only). IC_50_, defined as half maximal inhibitory concentration values and AUC (Area Under the Curve – %.mol.L^-1^) were obtained using library(ic50), library(drc), library(drda), library(ggplot2) and library (PharmacoGx) packages from R studio. Compounds were considered as potentially synergistic with the knockdown of *IRAK1* or the pharmacological inhibition of IRAK1 when the difference in AUC between the *IRAK1* silencing or the IRAK inhibitors and control conditions is more than 10%.mol.L^−1^. This cut-off corresponds to more than 3 times the S.D of the AUC of each library compound (< 3%) and considered as significant.

### Hit validation experiments

Following two-days transfection with siRNA targeted *IRAK1*, 4,000 living cells were seeded per well in 384-well plates (Corning, #3764). After 24h, cells were treated with a range of concentrations of BETi (molibresib and birabresib), MTAs (vincristine and vinorelbine), EGFRi (erlotinib and afatinib) and mTORi (everolimus and temsirolimus). Cell viability was measured using CellTiter-Glo^®^ 2.0 Cell Viability Assay (Promega, #G9243) after 72h of drug incubation in a humidified environment at 37°C and 5% CO_2_. Luminescence was measured using a PHERAstar^®^ plate reader (RRID:SCR_027001; BMG). Data were normalized to negative control wells (non-targeting negative control siRNA, DMSO). AUC were obtained using library(drda) package from R studio.

### Synergy experiments

Cells were seeded in 384-well U bottom and low-binding plates (#3830, Corning, USA). After 24h, cells were treated with a 6×5 combination matrix containing two compounds at different concentration ratios. After 7 days of drug treatment in a humidified environment at 37°C and 5% CO_2_, cell viability was measured using CellTiter-Glo^®^ 2.0 Cell Viability Assay (Promega, #G9243). Luminescence was measured using a PHERAstar^®^ plate reader (RRID:SCR_027001; BMG). Cell viability was determined and expressed as a percentage of untreated control cells. To assess the synergy score of the tested combinatorial treatments, Bliss heatmaps were generated using the SynergyFinder script in R studio ^25^. Images of the NB spheroids were acquired by phase contrast using a 4x/0.1 objective with a Leica DMIL LED^®^ (Leica) at the end point of experiments.

### NF-κB assays

Cells (3,000 BE(2)C, 4,000 SH-SY5Y per well) were seeded in 384-well plates (#3764, Corning). Twenty-four hours later, cells were transiently transfected using Lipofectamine 3000 (#L300015, Thermo Fisher Scientific) with the NF-κB - responsive NanoLuc reporter plasmid pNL3.2 NF-κB-RE [NlucP/NF-κB-RE/Hygro] and the constitutively expressed firefly luciferase control plasmid pGL4.54 [luc2/TK] (Promega; #N1111, #E5061). Forty-eight hours post-transfection, cells were pre-treated for 1h with recombinant human TNFα (20 ng/mL; R&D Systems) or DMSO, followed by treatment with IRAK1/4 inhibitor (1 µM), vincristine (10 nM), or a combination of both for 5h. NF-κB transcriptional activity was assessed 6h after stimulation using the Nano-Glo^®^ Dual-Luciferase^®^ Reporter Assay System (#N1620, Promega), according to the manufacturer’s instructions. Chemiluminescence was measured using a Pherastar microplate reader (BMG Labtech), and NF-κB activity was quantified as the ratio of NanoLuc to firefly luciferase signals.

### Cell cycle analysis and apoptosis detection

For cell cycle analysis, adherent and floating cells were harvested, fixed in 70% ethanol and incubated 15 min in the dark with 1 µL of FxCycle™ Violet stain, according to the manufacturer’s instructions (#F10347; Thermo-Fisher). The amount of DNA present was measured by flow cytometry (FACSymphony A3 analyzer; RRID:SCR_023644; BD Biosciences). For apoptosis detection, adherent and floating cells were harvested and incubated for 15 min in the dark with PI and Annexin V-FITC (#V13245; Thermo-Fisher), followed immediately by flow cytometry analysis (FACSymphony A3 analyzer; RRID:SCR_023644; BD Biosciences). Cytogram analysis was done with FlowJo software (RRID:SCR_008520; Tree Star, Ashland, OR, USA).

### scRNA-seq experiment acquisition

Cells were seeded in T75 cell culture flasks and treated with the different conditions. Twenty-four hours post-treatment, cells were fixed following the CG000478 protocol (10X Genomics). Briefly, cells were pelleted, fixed with 4% formaldehyde solution during 18h at 4°C, quenched and then stored at -80°C. Fixed cells were then processed following the Chromium Fixed RNA Profiling protocol for multiplexed samples (CG000527) using mouse probes. One million cells were incubated with specific mouse probes over night at 42°C. Samples were pooled to have the same number of cells for each sample. Pooled cells were then encapsulated to target around 5,000 cells per sample. Library was then generated and sequenced on Illumina Novaseq 6000 sequencer (RRID:SCR_016387) to target 70,000 reads/cells. All scRNAseq experiments were performed at the Cancer Genomic Platform of the Cancer Research Center of Lyon (UMR INSERM 1052 / CNRS 5286 / UCBL1/Centre Léon Bérard).

### scRNA-seq analysis

Raw sequencing data were processed using the Cell Ranger pipeline (10x Genomics) to generate gene-by-cell count matrices, which were subsequently imported into R for downstream analysis using Seurat (RRID:SCR_007322, v.5.1.0) ^53^. Each sample was preprocessed individually. Genes detected in fewer than 20 cells were excluded from further analysis. Ambient RNA contamination was estimated and corrected using SoupX (RRID:SCR_019193) ^54^, with parameters tfidfMin=0.5 and soupQuantile=0.85. Doublets were identified and removed by intersecting predictions from scDblFinder (RRID:SCR_022700) ^55^ and the hybrid method implemented in the scds package ^56^. Low-quality cells were filtered out based on mitochondrial content (>10%) and low library complexity, defined as cells with fewer than three median absolute deviations below the median in both the number of detected features and total UMI counts. Data were normalized using sctransform (RRID:SCR_022146) with default parameters, retaining 6,000 highly variable genes and regressing out effects associated with sequencing depth and cell cycle, the latter computed using Seurat’s cell cycle scoring function. After quality control and normalization, samples were merged into a single object, and dimensionality reduction was performed using principal component analysis (50 PCs), followed by UMAP for visualization. Neighbor graph construction and clustering were carried out using Seurat’s FindNeighbors and FindClusters functions respectively, with a final resolution of 0.15 selected after systematic comparison of multiple resolutions. Differential gene expression analyses for all relevant comparisons were performed using the Wilcoxon rank-sum test as implemented in Seurat’s FindMarkers function, applying Bonferroni correction and a minimum expression threshold of min.pct = 0.05. Gene set enrichment analysis for all comparisons was conducted with package fgsea (RRID:SCR_020938) ^57^ using Gene Ontology Biological Processes, Reactome, Hallmark, and KEGG MEDICUS gene sets, restricting analyses to gene sets containing between 5 and 500 genes. Genes were ranked using a combined metric incorporating adjusted p-values and log2FC.

### Univariate gene expression analysis

Gene expression analysis was conducted using the R2 microarray analysis and visualization platform (http://r2.amc.nl). Three independent cohorts providing open access to RNA-seq data were used including the SEQC and the Kocak databases for NB patient cohorts and the Genotype-Tissue Expression (GTeX) for normal adrenal gland dataset. Median values of the top 195 identified genes and *IRAK1*, *IRAK2*, *IRAK3* and *IRAK4* were recorded using log2 transformation gene expression. Statistical analyses using ANOVA were performed to compare IRAK isoform gene expression in NB patients to normal adrenal gland tissue gene expression. Boxplots were generated using GraphPad Prism 10.3.0 (RRID: SCR_002798).

### Multivariate gene expression analysis

The inputs of this analysis belong to data generated from Neuroblastoma patients’ tumor through the SEQC dataset that can be retrieved *via* the GEO omnibus entry GSE62564. Our study focuses on *IRAK* isoforms expression and clinical variables including age at diagnosis, INSS tumour stage and *MYCN* status for assessing patient prognosis. Patients with incomplete/missing information are discarded to perform the analysis, thus 493 out of 498 patients remain after data cleaning. Survival analyses were performed using R (version 4.1.1) with the survival (RRID: SCR_021137, version 3.4.0) and survminer (RRID: SCR_021094, version 0.4.9) packages. Patients were stratified into *IRAK* isoforms high- and low-expression groups based on Maximally Selected Rank Statistics provided in survminer. Hazard ratios (HRs) and 95% confidence intervals (CIs) were estimated applying univariate and multivariate Cox proportional hazards regression models. An HR > 1 indicates an increased risk of death compared to the reference (according to the definition of the hazard function) if a specific condition is met by a patient. Covariates constituted by our clinical variables of interest were included in multivariate models where appropriate. Statistical differences between groups were evaluated using the likelihood ratio test and *p*-values below 0.05 were considered statistically significant.

### Patient-derived neuroblastoma tumoroid models

Three previously generated patient-derived tumoroid models were included in this study ^27^. Patient-derived xenograft models were provided by the St. Jude Children’s Research Hospital. All animal studies were performed in strict compliance with relevant guidelines validated by the local Animal Ethics Evaluation Committee (C2EA-15) and authorized by the French Ministry of Education and Research (Authorization APAFIS#28836). Human tissue sample was obtained through a biopsy performed at Centre Léon Bérard. This sample was collected in the context of patient diagnosis. The Biological Resource Centre of the Centre Léon Bérard (n°BB-0033-463 00050) and the biological material collection and retention activity are declared to the Ministry of Research (DC-2008-99 and AC-2019-3426). The study had all necessary regulatory approvals and informed consents were available for all patients. NB tumoroids were expanded in 96-well ULA plates (#7007, Corning, USA) as previously described ^27^. Medium was changed twice a week and NB tumoroids were split every 7-10 days when reaching a diameter of 500-600μm (PDX-NB1) and 800-1,000μm (PDX-NB2 and -NB3) using TrypLE Express Enzyme (ThermoFisher Scientific, #12605036). All cultures were tested monthly for mycoplasma using the MycoAlert Mycoplasma Detection Kit (MycoAlert® Assay LONZA #LT07-710 and #LT07-518).

### Drug sensitivity assessment on tumoroid models

NB tumoroids were dissociated into individual cells using TrypLE Express (Thermo Fisher Scientific, #12605036). The cells were seeded at a density of 10,000 (PDX-NB2 and NB3) or 15,000 (PDX-NB1) cells per well in 96-well ULA plates (#7007, Corning, USA). The NB tumoroids were treated after 4 days of culture for 72 hours. For the cytotoxicity assay, the percentage of cell death was measured using a CellTox™ Green Cytotoxicity Assay (Promega, #G8743), according to the manufacturer’s instructions. All acquisitions were performed on a microplate reader (Tecan SPARK Multimode Microplate Reader (RRID:SCR_021897)). Data were normalized using negative (untreated controls) and positive (Triton X-100, 0.005%) control wells. Dose-response curves were obtained using GraphPad Prism 10.3.0 (RRID: SCR_002798). For the phenotypic LIVE/DEAD assays, the NB tumoroids were stained using the LIVE/DEAD**™** Viability/Cytotoxicity Kit (Thermo Fisher Scientific, #10237012) according to the manufacturer’s protocol, then imaged using a Leica Thunder Imager Live Cell system (RRID:SCR_023794). Immunofluorescence staining was automatically quantified by measuring the surface area of live (green) and dead (red) cells on at least 20 tumoroids using Fiji (RRID:SCR_002285) and then plotted as a ratio using GraphPad Prism 10.3.0 (RRID: SCR_002798).

### Tumoroid regrowth experiment

Three days after treatment, the NB tumoroids were washed 3 times in Advanced DMEM/F-12 (Gibco; #12491015) to remove the drugs, then placed in fresh complete culture medium. The culture medium was renewed twice a week and regrowth was carefully monitored using Cytena CELLCYTE X (RRID:SCR_021911). Once the maximal sustainable size for each NB tumoroid was reached in non-treated controls (800-1,000μm for PDX-NB2 and NB3), the experiment was stopped. All statistical analyses were performed on at least 10 organoids using GraphPad Prism 10.3.0 (RRID: SCR_002798).

### Animal study

*In vivo* NB experiments are approved under the Technion animal experimentation protocol No: IL-067-03-23H. Briefly, 8-week-old C57/BL6 mice were obtained from Envigo CRS (Israel). Mice were orthotopically injected with 0.5 million 9464D cells, a *MYCN-driven NB cell line* derived from TH-*MYCN* transgenic mice suspended in 10µL HBSS into the left adrenal gland. For the orthotopic injection, we performed microsurgery, using small incision to reach the adrenal gland, then cancer cell injection took place using Hamilton syringe. Mice were sedated with isoflurane 3% delivered through cannula. Following the surgery, analgesic was given - buprenorphine in dose of 0.05 mg/kg was administered by intraperitoneal injection. Experimental treatments started from day 5 post implantation and given every 3 days until endpoint. Vincristine was dissolved in sterile PBS while IRAK1/4i in corn oil as well as DMSO for the vehicle condition. All treatments were administered by intraperitoneal injection with a final volume of 100µL at the following concentrations: 0.4 mg/kg and 7.5 mg/kg for vincristine and IRAK1/4i respectively. Mice (n ≥ 5/group) were clinically assessed twice a week including behavior evaluation, clinical assessment score and weight. Tumor size was determined twice a week using micro ultra-sounds (VisualSonics Vevo^®^ 3100 Imaging system; RRID:SCR_022921). All images were recorded and processed in an *in vivo* analysis software Vevo LAB. Tumor volume was calculated according to the formula ½ x width x depth x length and expressed in mm^3^. Mice were allowed to live until their natural death or were sacrificed when their tumor volume exceeded 1,000mm^3^ or their weight decrease in more than 20%. Survival medians were estimated by the Kaplan-Meier product limit method. The log-rank test was used to compare survival rates by univariate analysis.

### Statistical analysis

Sample size and replicates are stated in the corresponding figure legends. Data are presented as mean ± S.D. Statistical significance was tested using unpaired Student’s t test. For experiments using multiple variables, statistical significance was assessed *via* two-way ANOVA. A significant difference between two conditions was recorded for \**p* < 0.05; \*\**p* < 0.01; \*\*\**p* < 0.001.

## ACKNOWLEDGEMENTS

We thank Dr Xavier Morelli and Carine Derviaux of the HiTS drug screening platform of CRCM for their assistance with the functional and pharmacological drug screens. We would like to thank all of our funding sources. This work was exclusively funded by institutional grants and donations from non-for-profit organizations, which had no influence on the direction of this research. It included research grants attributed to EP or MLG: “Generic call” grant from the AMIDEX of Aix Marseille University, post-doctoral fellowship from the ARC Foundation (#ARCPDF22020070002553), “Emergence” grant from the Canceropole PACA and research grants from the not-for-profit organizations Association Cassandra, Wonder Augustine and Eva pour la Vie. This work was also supported by the PEDIACRIEX grant from the French National Cancer Institute for the creation of the Lyon-Marseille Centre of Excellence in Pediatric Oncology Research, South-ROCK (INCa-Cancer_18695), La Ligue Nationale Contre le Cancer (Equipes labellisées 2025) and by donations from several not-for-profit organizations (Association Léa, Le sourire de Lucie, Vocaliz’ et Nos p’tites Etoiles).

## AUTHOR CONTRIBUTIONS

Conceptualization: MLG and EP

Methodology: MLG, CB, AMR, KM, AB, TWF, GMA, FMGC, LB and EP

Investigation: MLG, CB, MK, FG, AMR, MT, KM, BM, SC, AB, SL, EL and EP

Funding acquisition: MLG and EP

Project administration: MLG and EP

Supervision: MLG, YS, LB, FMGC, NA and EP

Providing materials: MLG, CB, FG, LB, YS and EP

Writing – original draft: MLG, CB, AMR, AB, LB and EP

Writing – review & editing: MLG, CB, MK, MT, AMR, FMGC, NA, YS and EP.

## REFERENCES

1. Scott, E. C. et al. Trends in the approval of cancer therapies by the FDA in the twenty-first century. Nat. Rev. Drug Discov. 22, 625–640 (2023).

2. Hopkins, A. L. Network pharmacology: the next paradigm in drug discovery. Nat. Chem. Biol. 4, 682–690 (2008).

3. Al-Lazikani, B., Banerji, U. & Workman, P. Combinatorial drug therapy for cancer in the post-genomic era. Nat. Biotechnol. 30, 679–692 (2012).

4. Wishart, D. S. et al. DrugBank 5.0: a major update to the DrugBank database for 2018. Nucleic Acids Res. 46, D1074–D1082 (2018).

5. Amelio, I. et al. DRUGSURV: a resource for repositioning of approved and experimental drugs in oncology based on patient survival information. Cell Death Dis. 5, e1051 (2014).

6. Gaulton, A. et al. The ChEMBL database in 2017. Nucleic Acids Res. 45, D945–D954 (2017).

7. Peters, J.-U. Polypharmacology - foe or friend? J. Med. Chem. 56, 8955–8971 (2013).

8. Anighoro, A., Bajorath, J. & Rastelli, G. Polypharmacology: challenges and opportunities in drug discovery. J. Med. Chem. 57, 7874–7887 (2014).

9. Zhang, W., Bai, Y., Wang, Y. & Xiao, W. Polypharmacology in Drug Discovery: A Review from Systems Pharmacology Perspective. Curr. Pharm. Des. 22, 3171–3181 (2016).

10. Ariey-Bonnet, J. et al. Combination drug screen targeting glioblastoma core vulnerabilities reveals pharmacological synergisms. eBioMedicine 95, 104752 (2023).

11. Maris, J. M. Recent advances in neuroblastoma. N. Engl. J. Med. 362, 2202–2211 (2010).

12. Irwin, M. S. & Park, J. R. Neuroblastoma: paradigm for precision medicine. Pediatr. Clin. North Am. 62, 225–256 (2015).

13. Qiu, B. & Matthay, K. K. Advancing therapy for neuroblastoma. Nat. Rev. Clin. Oncol. 19, 515–533 (2022).

14. Oesterheld, J. et al. Eflornithine as Postimmunotherapy Maintenance in High-Risk Neuroblastoma: Externally Controlled, Propensity Score–Matched Survival Outcome Comparisons. J. Clin. Oncol. 42, 90–102 (2024).

15. Pastor, E. R. & Mousa, S. A. Current management of neuroblastoma and future direction. Crit. Rev. Oncol. Hematol. 138, 38–43 (2019).

16. Su, L.-C., Xu, W.-D. & Huang, A.-F. IRAK family in inflammatory autoimmune diseases. Autoimmun. Rev. 19, 102461 (2020).

17. Wee, Z. N. et al. IRAK1 is a therapeutic target that drives breast cancer metastasis and resistance to paclitaxel. Nat. Commun. 6, 8746 (2015).

18. Zhang, D. et al. Constitutive IRAK4 Activation Underlies Poor Prognosis and Chemoresistance in Pancreatic Ductal Adenocarcinoma. Clin. Cancer Res. Off. J. Am. Assoc. Cancer Res. 23, 1748–1759 (2017).

19. Cheng, B. Y. et al. IRAK1 Augments Cancer Stemness and Drug Resistance via the AP-1/AKR1B10 Signaling Cascade in Hepatocellular Carcinoma. Cancer Res. 78, 2332–2342 (2018).

20. Liu, P. H. et al. An IRAK1-PIN1 signalling axis drives intrinsic tumour resistance to radiation therapy. Nat. Cell Biol. 21, 203–213 (2019).

21. Liu, L. et al. Targeting the IRAK1-S100A9 Axis Overcomes Resistance to Paclitaxel in Nasopharyngeal Carcinoma. Cancer Res. 81, 1413–1425 (2021).

22. Kim, K. M., Hwang, N.-H., Hyun, J.-S. & Shin, D. Recent Advances in IRAK1: Pharmacological and Therapeutic Aspects. Molecules 29, 2226 (2024).

23. Cotto, K. C. et al. DGIdb 3.0: a redesign and expansion of the drug-gene interaction database. Nucleic Acids Res. 46, D1068–D1073 (2018).

24. Rhyasen, G. W. & Starczynowski, D. T. IRAK signalling in cancer. Br. J. Cancer 112, 232–237 (2015).

25. Ianevski, A., Giri, A. K. & Aittokallio, T. SynergyFinder 3.0: an interactive analysis and consensus interpretation of multi-drug synergies across multiple samples. Nucleic Acids Res. gkac382 (2022) doi:10.1093/nar/gkac382.

26. Barbet, V. & Broutier, L. Future Match Making: When Pediatric Oncology Meets Organoid Technology. Front. Cell Dev. Biol. 9, 674219 (2021).

27. Ma, X. et al. MYC shapes ER-mitochondria calcium transfer by directly targeting *ITPR1* : implications for MYC-induced safeguard mechanisms and cancer. Preprint at 10.1101/2024.08.28.610025 (2024).

28. Li, J. et al. Radiation induces IRAK1 expression to promote radioresistance by suppressing autophagic cell death via decreasing the ubiquitination of PRDX1 in glioma cells. Cell Death Dis. 14, 259 (2023).

29. Wong, M. et al. Whole genome, transcriptome and methylome profiling enhances actionable target discovery in high-risk pediatric cancer. Nat. Med. 26, 1742–1753 (2020).

30. Van Tilburg, C. M. et al. The Pediatric Precision Oncology INFORM Registry: Clinical Outcome and Benefit for Patients with Very High-Evidence Targets. Cancer Discov. 11, 2764–2779 (2021).

31. Berlanga, P. et al. The European MAPPYACTS Trial: Precision Medicine Program in Pediatric and Adolescent Patients with Recurrent Malignancies. Cancer Discov. 12, 1266–1281 (2022).

32. Molenaar, J. J. et al. Sequencing of neuroblastoma identifies chromothripsis and defects in neuritogenesis genes. Nature 483, 589–593 (2012).

33. Pugh, T. J. et al. The genetic landscape of high-risk neuroblastoma. Nat. Genet. 45, 279–284 (2013).

34. Nikolic, I. et al. Discovering cancer vulnerabilities using high-throughput micro-RNA screening. Nucleic Acids Res. 45, 12657–12670 (2017).

35. Soriano, A. et al. Functional high-throughput screening reveals miR-323a-5p and miR-342-5p as new tumor-suppressive microRNA for neuroblastoma. Cell. Mol. Life Sci. CMLS 76, 2231–2243 (2019).

36. Lochmann, T. L. et al. Targeted inhibition of histone H3K27 demethylation is effective in high-risk neuroblastoma. Sci. Transl. Med. 10, eaao4680 (2018).

37. Vernooij, L. et al. High-Throughput Screening Identifies Idasanutlin as a Resensitizing Drug for Venetoclax-Resistant Neuroblastoma Cells. Mol. Cancer Ther. 20, 1161–1172 (2021).

38. Zaghmi, A. et al. High-content screening of drug combinations of an mPGES-1 inhibitor in multicellular tumor spheroids leads to mechanistic insights into neuroblastoma chemoresistance. Mol. Oncol. 18, 317–335 (2023).

39. Herter, S. et al. High content-imaging drug synergy screening identifies specific senescence-related vulnerabilities of mesenchymal neuroblastomas. Cell Death Dis. 16, 644 (2025).

40. Pereira, M. & Gazzinelli, R. T. Regulation of innate immune signaling by IRAK proteins. Front. Immunol. 14, 1133354 (2023).

41. Aggarwal, R. K. et al. Smoking-Associated Carcinogen-Induced Inflammation Promotes Lung Carcinogenesis via IRAK4 Activation. Clin. Cancer Res. Off. J. Am. Assoc. Cancer Res. 31, 746–755 (2025).

42. Zhang, D. et al. Tumor-Stroma IL1β-IRAK4 Feedforward Circuitry Drives Tumor Fibrosis, Chemoresistance, and Poor Prognosis in Pancreatic Cancer. Cancer Res. 78, 1700–1712 (2018).

43. Liu, M. et al. A Pan-Cancer Analysis of IRAK1 Expression and Their Association With Immunotherapy Response. Front. Mol. Biosci. 9, 904959 (2022).

44. Wang, Y., Wang, Y., Duan, X., Wang, Y. & Zhang, Z. Interleukin-1 receptor-associated kinase 1 correlates with metastasis and invasion in endometrial carcinoma. J. Cell. Biochem. 119, 2545–2555 (2018).

45. Chen, W. et al. IRAK1 deficiency potentiates the efficacy of radiotherapy in repressing cervical cancer development. Cell. Signal. 119, 111192 (2024).

46. O’Neil, J. et al. An Unbiased Oncology Compound Screen to Identify Novel Combination Strategies. Mol. Cancer Ther. 15, 1155–1162 (2016).

47. Jaaks, P. et al. Effective drug combinations in breast, colon and pancreatic cancer cells. Nature 603, 166–173 (2022).

48. Li, Z. et al. Inhibition of IRAK1/4 sensitizes T cell acute lymphoblastic leukemia to chemotherapies. J. Clin. Invest. 125, 1081–1097 (2015).

49. Kim, K. M., Hwang, N.-H., Hyun, J.-S. & Shin, D. Recent Advances in IRAK1: Pharmacological and Therapeutic Aspects. Mol. Basel Switz. 29, 2226 (2024).

50. Li, Y. et al. A non-canonical IRAK4-IRAK1 pathway counters DNA damage-induced apoptosis independently of TLR/IL-1R signaling. Sci. Signal. 16, eadh3449 (2023).

51. Pasquier, E. et al. β-blockers increase response to chemotherapy via direct antitumour and anti-angiogenic mechanisms in neuroblastoma. Br. J. Cancer 108, 2485–2494 (2013).

52. Zhang, null, Chung, null & Oldenburg, null. A Simple Statistical Parameter for Use in Evaluation and Validation of High Throughput Screening Assays. J. Biomol. Screen. 4, 67–73 (1999).

53. Hao, Y. et al. Dictionary learning for integrative, multimodal and scalable single-cell analysis. Nat. Biotechnol. 42, 293–304 (2024).

54. Young, M. D. & Behjati, S. SoupX removes ambient RNA contamination from droplet-based single-cell RNA sequencing data. GigaScience 9, giaa151 (2020).

55. Germain, P.-L., Lun, A., Meixide, C. G., Macnair, W. & Robinson, M. D. Doublet identification in single-cell sequencing data using *scDblFinder*. Preprint at 10.12688/f1000research.73600.2 (2022).

56. Bais, A. S. & Kostka, D. scds: computational annotation of doublets in single-cell RNA sequencing data. Bioinformatics 36, 1150–1158 (2020).

57. Korotkevich, G. et al. Fast gene set enrichment analysis. 060012 Preprint at 10.1101/060012 (2021).

