## Supplementary material for "Reverse molecular pharmacology identifies the non-canonical axis of IRAK as a chemoresistance factor in neuroblastoma": Suppl Figure

### Supplementary Figures

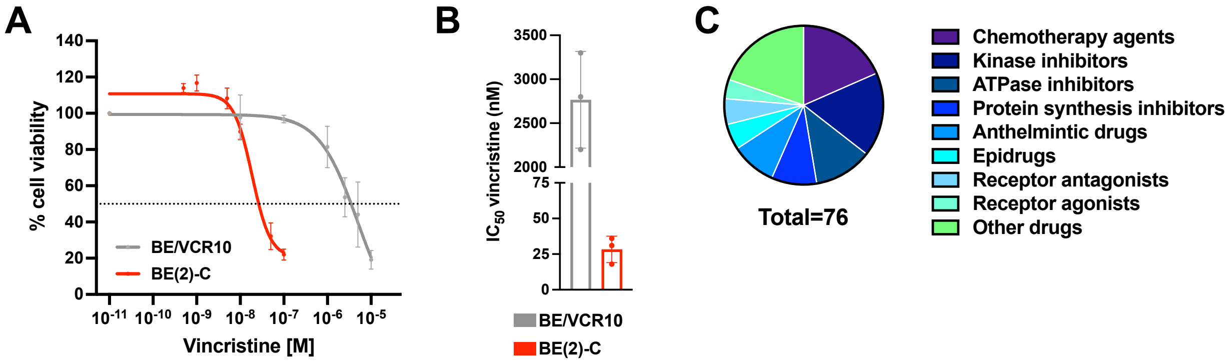


**Supplementary Figure 1:** (**A**) Cell survival assay performed on BE(2)-C cells and the BE/VCR10 resistant cell line after 72 h incubation with a range of concentrations of Vincristine. Means of at least three individual experiments: bars, S.D; log scale for x axis. (**B**) Histogram representation of the $IC_{50}$ values in BE(2)-C cells and the BE/VCR10 resistant cell line. Means of three individual experiments: bars, S.D. (**C**) Seventy-six compounds were defined as NB killers and are represented in donut diagram and classified by pharmacological classes.

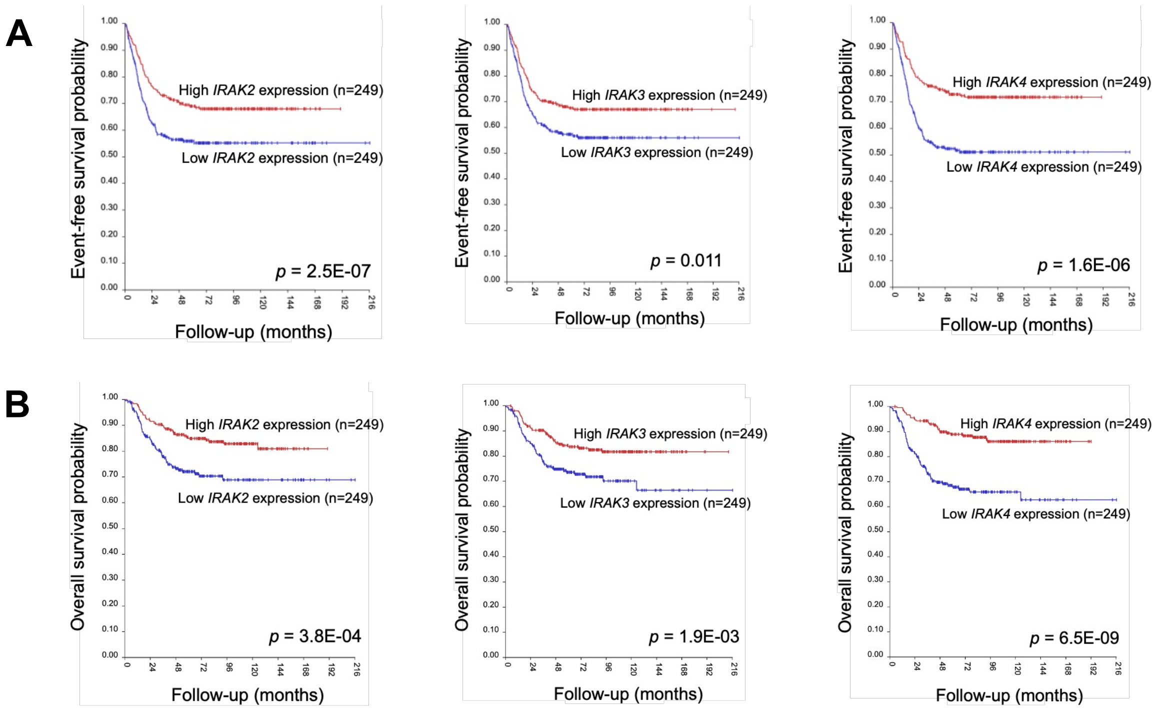


**Supplementary Figure 2:** (**A, B**) Kaplan–Meier curves showing the probability of EFS and OS for the SEQC cohort. Values were dichotomized into 'high' and 'low' *IRAK isoform* expression around the median.

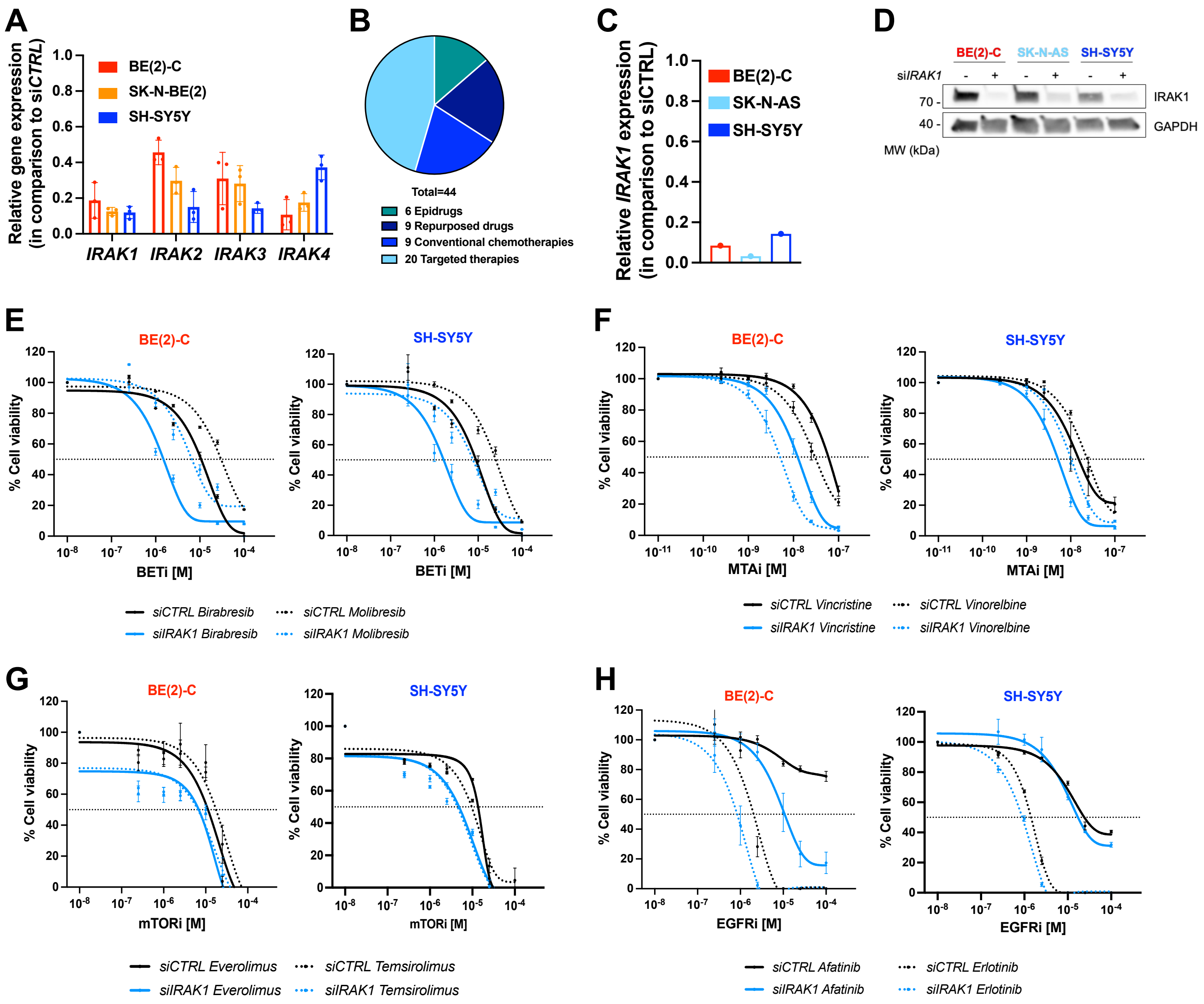


**Supplementary Figure 3:**(**A**) Relative gene expression following 48h transfection of BE(2)-C, SK-N-BE(2) and SH-SY5Y cells with negative control siRNA and siRNA sequences targeting the 4 IRAK isoforms, as evaluated by qRT-PCR using *YWHAZ* as housekeeping gene. (**B**) Forty-four compounds were used for the functional drug screening and are represented in donut diagram and classified by pharmacological classes. (**C**) Relative gene expression following 48h transfection of BE(2)-C, SK-N-AS and SH-SY5Y cells with negative control siRNA and siRNA sequence targeting *IRAK1*, as evaluated by qRT-PCR using *YWHAZ* as housekeeping gene. (**D**) Representative western blot showing IRAK1 protein expression following 72h siRNA transfection in BE(2)-C, SK-N-AS and SH-SY5Y cells, using GAPDH as loading control. (**E,F,G,H**) Dose–response curves of BE(2)-C or SH-SY5Y cells either transfected with negative control or siRNA targeting *IRAK1*, treated with (**E**) BETi, (**F**) MTAs, (**G**) mTORi and (**H**) EGFRi for 72 hours.

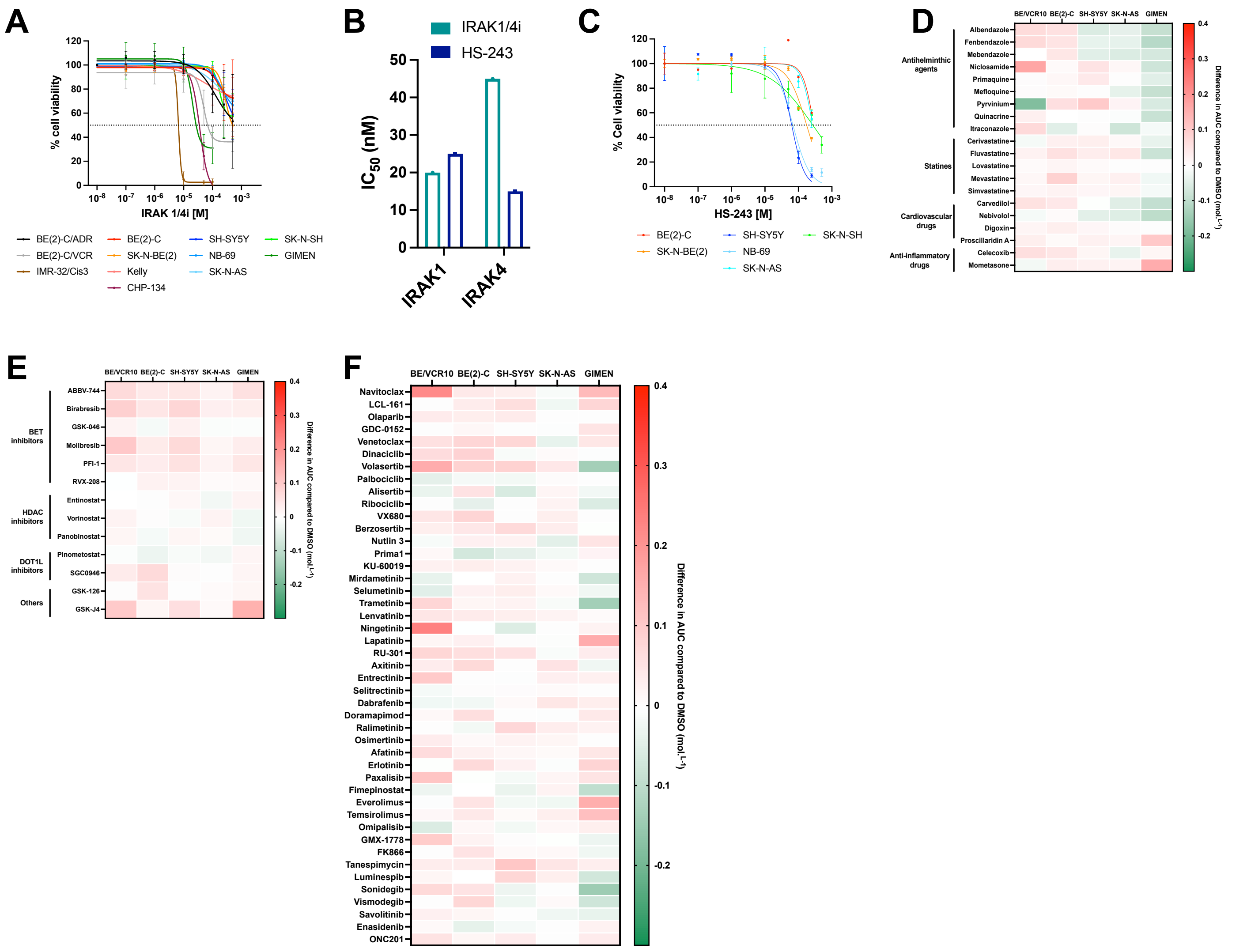


**Supplementary Figure 4:** (**A**) Dose–response curves of 12 NB cells treated with IRAK1/4i for 72 hours. (**B**) Histograms representing the $IC_{50}$ values of IRAK1/4i and HS-243 for IRAK1 or IRAK4 calculated with a kinase assay. (**C**) Dose–response curves of 6 NB cells treated with HS-243 for 72 hours. (**D, E, F**) Heat maps classification representing the differences in AUC between the average of both IRAK inhibitors and DMSO conditions in BE/VCR10, BE(2)-C, SH-SY5Y, SK-N-AS and GIMEN cells lines for the (**D**) 20 repurposed drugs, (**F**) 13 epidrugs and (**E**) 45 targeted therapies.

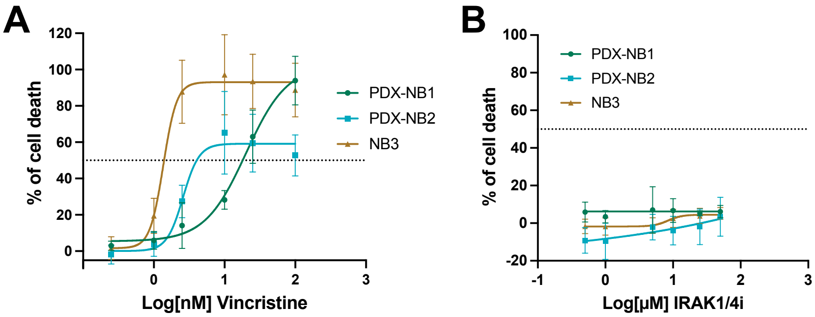


**Supplementary Figure 5: (A-B)** Dose-response curves showing cell death (%) in PDX-NB1, PDX-NB2, and NB3 organoids treated with vincristine **(A)** or IRAK1/4i **(B)** for 72h. Cell death is expressed as percentage relative to CellTox-Green-treated control (TritonX100; 0,005%). Values represent the mean of at least three independent experiments ± S.D.

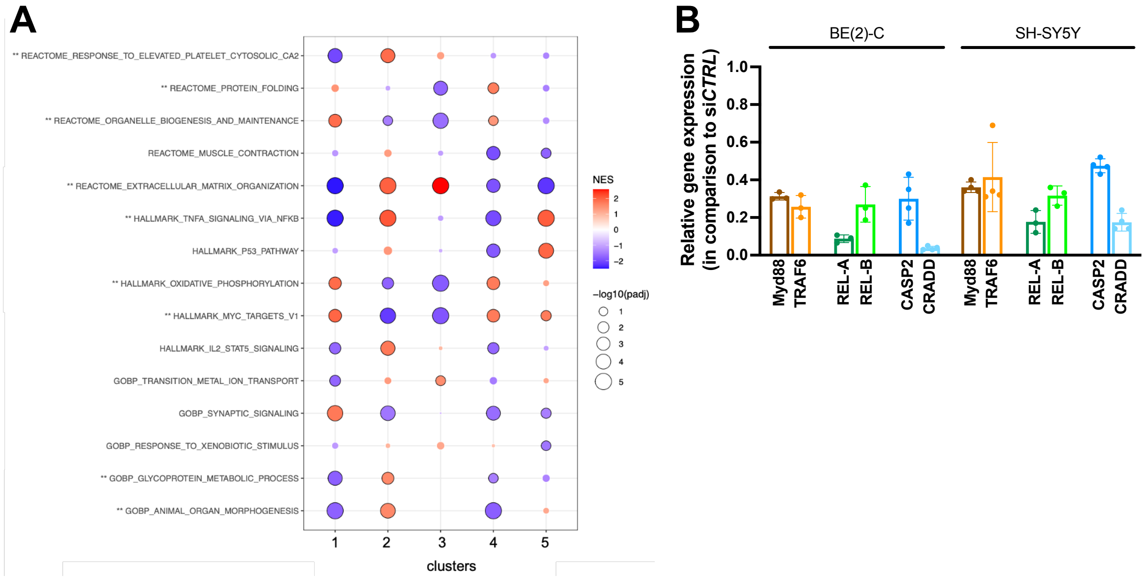


**Supplementary Figure 6: (A)** Dot plot representing gene expressions between cluster groups. (**B**) Relative gene expression following 48h transfection of BE(2)-C and SH-SY5Y cells with negative control siRNA and siRNA sequence targeting *Myd88, TRAF6, REL-A,REL-B, CASP2* or *CRADD* as evaluated by qRT-PCR using *YWHAZ* as housekeeping gene. Values are the average of at least three independent experiments ± S.D; ns, p>0.05.
