## Supplementary material for "Reverse molecular pharmacology identifies the non-canonical axis of IRAK as a chemoresistance factor in neuroblastoma": Suppl Tables 5 and 6

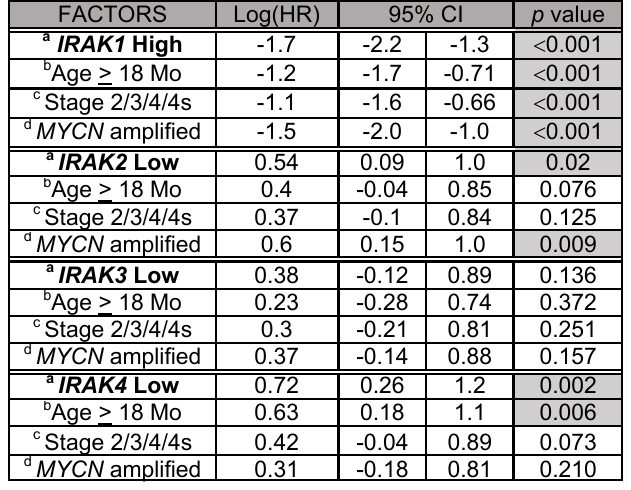

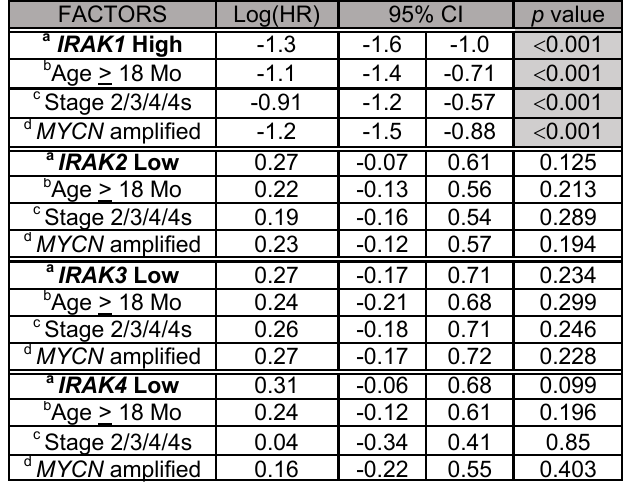

**Supplementary Table 6: Multivariate analysis of overall survival (OS)**

**of neuroblastoma patients (n=498, SEQC cohort)**

^a^ *^IRAK^* ^isoform gene expression (maximally selected rank statistics applied for optimal cut-points)^

^b Age at diagnosis ≥18 months^

^c Tumor determined to be INSS stage 2, 3 4 or 4s at diagnosis^

^d Amplification of^ *^MYCN^* ^gene^

HR – Hazard Ratio. An HR > 1 indicates a poor prognosis compared to the reference. CI – Confidence Interval

**Supplementary Table 5: Multivariate analysis of event-free survival (EFS) of neuroblastoma patients (n=498, SEQC cohort)**
